# Ocean currents drive the worldwide colonization of the most widespread marine plant, eelgrass (*Zostera marina*)

**DOI:** 10.1101/2022.12.10.519859

**Authors:** Lei Yu, Marina Khachaturyan, Michael Matschiner, Adam Healey, Diane Bauer, Brenda Cameron, Mathieu Cusson, J. Emmet Duffy, F. Joel Fodrie, Diana Gill, Jane Grimwood, Masakazu Hori, Kevin Hovel, A. Randall Hughes, Marlene Jahnke, Jerry Jenkins, Keykhosrow Keymanesh, Claudia Kruschel, Sujan Mamidi, Per-Olav Moksnes, Masahiro Nakaoka, Christa Pennacchio, Katrin Reiss, Francesca Rossi, Jennifer L. Ruesink, Stewart Schultz, Sandra Talbot, Richard Unsworth, Tal Dagan, Jeremy Schmutz, John J. Stachowicz, Yves Van de Peer, Jeanine L. Olsen, Thorsten B. H. Reusch

## Abstract

Currents are unique drivers of oceanic phylogeography and so determine the distribution of marine coastal species, along with past glaciations and sea level changes. Here, we reconstruct the worldwide colonization history of eelgrass (*Zostera marina* L.), the most widely distributed marine flowering plant or seagrass from its origin in the Northwest Pacific, based on nuclear and chloroplast genomes. We identified two divergent Pacific clades with evidence for admixture along the East Pacific coast. Multiple west to east (trans-Pacific) colonization events support the key role of the North Pacific Current. Time-calibrated nuclear and chloroplast phylogenies yielded concordant estimates of the arrival of *Z. marina* in the Atlantic through the Canadian Arctic, suggesting that eelgrass-based ecosystems, hotspots of biodiversity and carbon sequestration, have only been present since ∼208 Kya (thousand years ago). Mediterranean populations were founded ∼53 Kya while extant distributions along western and eastern Atlantic shores coincide with the end of the Last Glacial Maximum (∼20 Kya). The recent colonization and 5-to 7-fold lower genomic diversity of Atlantic compared to the Pacific populations raises concern and opportunity about how Atlantic eelgrass might respond to rapidly warming coastal oceans.

---

Seagrasses are the only flowering plants that returned to the sea ∼67 mya (million years ago), comprising at least three independent lineages that descended from freshwater ancestors ∼114 mya^1^. Seagrasses are foundation species of entire ecosystems thriving in all shallow coastal areas of the global ocean except Antarctica^2^. By far the most geographically widespread species is eelgrass (*Zostera marina*), occurring in Pacific and Atlantic areas of the northern hemisphere from warm temperate to Arctic environments^3^, spanning 40° of latitude and a range of ∼18°C in average annual temperatures (Fig. 1a). Eelgrass is a unique foundation species in that no other current seagrass can fill its ecological niche in the cold temperate to Arctic northern hemisphere^3^ (Supplementary Note 1).

**Fig. 1.**
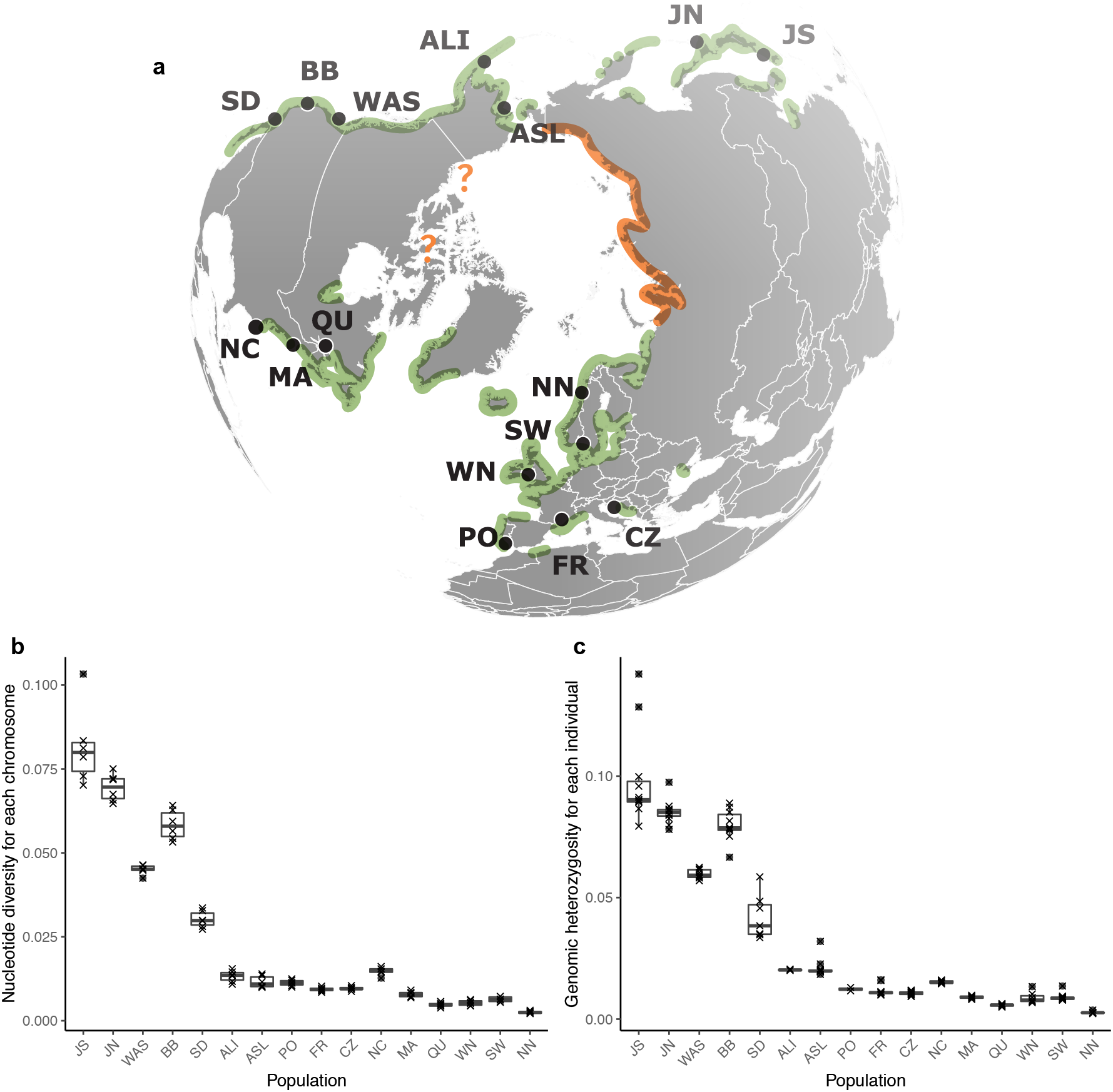
Distribution and sampling sites for *Zostera marina* and their widely varying genetic diversity. **a**, Population abbreviations: San Diego, California (SD); Bodega Bay, California (BB); Washington state (WAS); Alaska-Izembek (ALI); Alaska-Safety Lagoon (ASL); Japan-North (JN); Japan-South (JS); North Carolina (NC); Massachusetts (MA); Quebec (QU); Northern Norway (NN); Sweden (SW); Wales North (WN); Portugal (PO); Mediterranean France (FR); Croatia (CZ). Green areas indicate presence of *Z. marina*. The orange line along the Siberian coastline represents the absence of *Z. marina* based on cursory surveys of Alismatales including *Z. marina* by Russian colleagues. The latter areas are characterized by gravel coasts, river outflows and turbid waters. Question marks indicate areas that have not been explored. Detailed location metadata can be found in Supplementary Table 1. **b**, Genetic diversity: box-plots (25/75% percentile, median) of nucleotide diversity (π), calculated for each of the six chromosomes based on the 44,865 SNP set (Supplementary Fig. 1). Each data point indicates one chromosome. **c**, Box-plots of individual genome wide heterozygosity *H*_obs_ based on the 144,773 SNP set (Supplementary Fig. 1), as (number of heterozygous sites) / (total number of sites with genotype calls). Each data point corresponds to a population sample (10-12 individuals). See statistical tests for differences in mean π or *H*_obs_ in Supplementary Table 4.

Given its very wide natural distribution range that exceeds most terrestrial plant species, our goal was to reconstruct the major colonization pathways of eelgrass starting from its putative origin of *Z. marina* in the West Pacific along the Japanese Archipelago^4,5^. Currents are unique to phylogeographic processes in the ocean and we hypothesized that major current systems such as North Pacific and California Currents in the Pacific, and the Gulf Stream and North Atlantic Drift in the Atlantic drove its worldwide colonization.

To put data into perspective with rates of colonization in terrestrial plant species, one major goal was to provide time estimates of major colonization events. We asked specifically how evolutionary contingency—specifically large-scale dispersal events—may have affected the timing of arrival of eelgrass on East Pacific and North Atlantic coastlines^6^. To do so, we took advantage of recent extensions of the multispecies coalescent (MSC) as applied at the population level^7,8^, making it possible to construct a time-calibrated phylogenetic tree from SNP (single nucleotide polymorphism) data^9^. Our data set comprised 190 individuals from 16 worldwide locations that were subjected to comprehensive whole-genome resequencing (nuclear and chloroplast).

Superimposed onto the general eastward colonization are Pleistocene cycles of glacial and interglacial periods that resulted in frequent latitudinal expansions and contractions of available habitat for both terrestrial and marine biota^10^. Such local extinctions and subsequent recolonizations from refugial populations are expected to leave their genomic footprint in extant marine populations^11-13^ and may restrict their potential to rapidly adapt to current environmental change^14,15^. Hence, we were also interested in how glaciations—in particular the Last Glacial Maximum (LGM; 20,000 yrs ago (Kya)^16^)—have affected population-wide genomic diversity of *Z. marina*, and which glacial refugia permitted eelgrass to survive this period.

## Results

### Whole-genome resequencing and nuclear and chloroplast polymorphism

Among 190 *Z. marina* specimen collected from 16 geographic locations (Fig. 1a, Supplementary Table 1), full genome sequencing yielded an average read coverage of 53.73x. After quality filtering (Supplementary Data 1), single nucleotide polymorphisms (SNPs) were mapped and called (Supplementary Fig. 1,2) based on a chromosomal level assembly v.3.1^17^. In order to facilitate phylogenetic construction within a conserved set of genes^18^, we first assessed the presence of a core gene set shared by all individuals. From a total of 21,483 genes, we identified 18,717 core genes that were on average observed in 97% of samples, containing 763,580 SNPs (Supplementary Note 3).

After exclusion of 37 samples owing to missing data, selfing or clonality, 153 were left for further analyses (Supplementary Tables 2,3; Supplementary Fig. 3,4). We also obtained a thinned synonymous data set retaining only sites with a physical distance of >3 kbp (11,705 SNPs, hereafter “ZM_Core_SNPs”) (Supplementary Fig. 1,2).

A complete chloroplast genome of 143,968 bp was reconstructed from the reference sample^19^. Median chloroplast sequencing coverage for the samples of the worldwide data set was 6273x. A total of 151 SNPs were detected along the whole chloroplast genome, excluding 23S and 16S gene regions due to possible contamination in some samples and ambiguous calling next to microsatellite regions (132,438 bp), comprising 54 haplotypes.

### Gradients of genetic diversity within and among ocean basins

As measures of genetic diversity, we assessed nucleotide diversity (π) and genome-wide heterozygosity (*H*_obs_) (Fig. 1b,c). Consistent with the assumed Pacific origin of the species, Pacific locations displayed a 5.5 (π)-to 6.6 (*H*_obs_)-fold higher genetic diversity compared to the Atlantic (Supplementary Table 4). The highest π- and *H*_obs_-values were observed in Japan South (JS) followed by Japan North (JN), suggesting the origin of *Z. marina* in the Northwest Pacific^4,5^. Alaska Izembek (ALI) and Alaska Safety Lagoon (ASL) displayed approximately a third (28% for π; 34% for *H*_obs_) of the diversity in the more southern Pacific sites (average of San Diego SD, Bodega Bay BB, Washington State WAS). In the Atlantic, a comparable loss of diversity along a south-north gradient was observed. Quebec (QU) displayed 42% (π) and 47% (*H*_obs_) of the diversity of North Carolina (NC) and Massachusetts (MA), while the diversity values in Norway (NN) was 31% and 43% of averaged values of Sweden (SW) and Wales (WN).

### Global population structure of *Z. marina*

To reveal the large-scale population genetic structure, we performed a Principal Component Analysis (PCA) based on the most comprehensive SNP selection (Supplementary Fig. 1; 782,652 SNPs, Fig. 2a). Within-ocean genetic differentiation in the Pacific was as great as the Pacific-Atlantic split, whereas there was much less variation within the Atlantic. Separate PCAs for each ocean revealed additional structure (Fig. 2c,e), including the separation of the Atlantic and Mediterranean Sea populations (PC1, 24.47%, Fig. 2e).

**Fig. 2.**
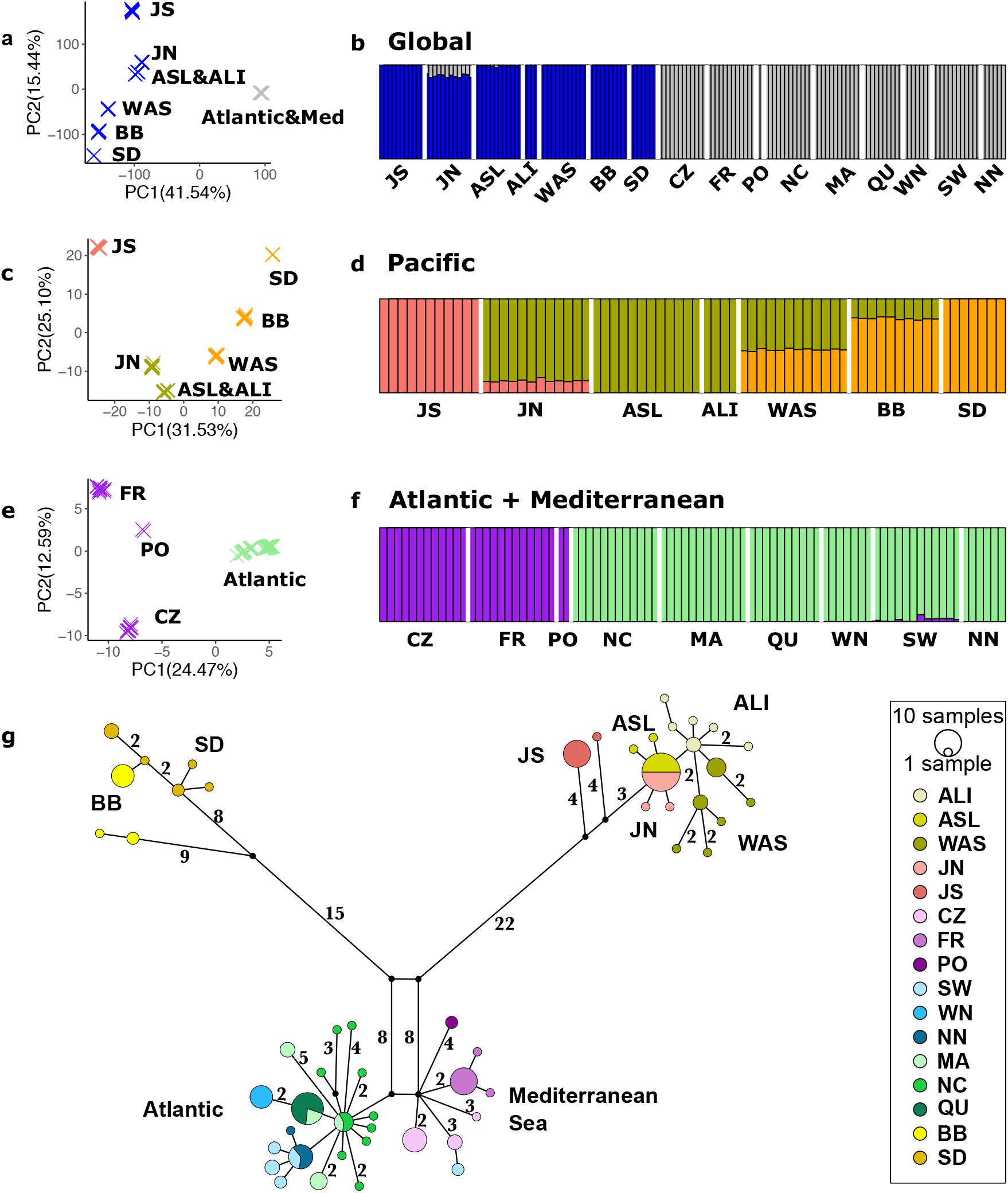
Population structure based on nuclear and cpDNA SNPs among 16 eelgrass populations. **a**,**b**, Global genetic population structure. **a**, Global Principal Component Analysis (PCA) based on 782,652 SNPs, here Atlantic and Mediterranean populations are collapsed. Pacific and Atlantic Ocean were separated by PC1 that explained 41.75% of the variation **b**, Global STRUCTURE analysis (no of clusters, K = 2; based on 2,353 SNPs). Each individual is represented by a vertical bar partitioned into colors based on its affiliation to a genetic cluster, as determined by delta-K method (see Methods) **c, d**, Genetic population structure within the Pacific. **c**, PCA within the Pacific based on 12,514 SNPs. **d**, STRUCTURE analysis within the Pacific (K = 3; 6,168 SNPs). **e, f**, Genetic population structure for the Atlantic and the Mediterranean Sea. **e**, PCA for the Atlantic and the Mediterranean Sea based on 8,552 SNPs. **f**, STRUCTURE analysis for the Atlantic and the Mediterranean Sea (K = 2; 8,552 SNPs). See Supplementary Fig. 5-7 for results assuming higher numbers of clusters, and Supplementary Fig. 1 for further details on the SNP sets used. **g**, cpDNA haplotype network. Numbers represent mutation steps >1. Colors correspond to the population. Split-colored circles indicate that a particular haplotype is shared between populations, circle size is proportional to frequency.

We then used STRUCTURE^20^, a Bayesian clustering approach, on 2,353 SNPs (20%) randomly selected from the ZM_Core_SNPs. The optimal number of genetic clusters was determined using the Delta-K method^21^ (Fig. 2b,d,f), with additional K-values explored in Supplementary Fig. 5-7. In the global analysis, (Fig. 2b), two clusters representing Atlantic and Pacific locations were identified. JN contained admixture components with the Atlantic, consistent with a West-East colonization via northern Japan through the North Pacific Current and then north towards the Bering Sea. An analysis restricted to Pacific sites (K=3) supported a role of JN as dispersal hub, with admixture components from JS and Alaska, suggesting that this site has been a gateway between both locations (Fig. 2c). WAS and BB, located centrally along the east Pacific coastline, were admixed between both Alaskan sites and SD. WAS displayed about equal northern and southern components, while BB was dominated by the adjacent southern SD genetic component. In the Atlantic (Fig. 2f), a less pronounced population structure was present, consistent with the PCA results (Fig. 2e). The optimal number of genetic clusters was K=2, separating the northern Atlantic and the Mediterranean, yet analyses with K=4 revealed a connection between Portugal (PO) closest to the Strait of Gibraltar and the East Atlantic (NC, Supplementary Fig. 7).

### Population structure of cpDNA

A haplotype network (Fig. 2g) revealed three markedly divergent clades. In the Pacific, WAS displayed haplotypes similar to those of Alaska (ALI/ASL) and JN, while BB displayed haplotypes of a divergent clade that also comprises all haplotypes from SD. ASL and JN share the same dominant haplotype, suggesting JN to be a hub between West and East Pacific respectively Alaska. In JS, two divergent private haplotypes (separated by nine mutations from other haplotypes) suggest long-term persistence of eelgrass at that location.

On the Atlantic side, only four to six mutations separate the Northeast Atlantic and Mediterranean haplotypes, consistent with a much younger separation. The central haplotype is shared by both MA and NC, with nine private NC haplotypes. A single mutation separates both MA and QU; and MA and WN. Also extending from the central haplotype were SW and NN. Together with the diversity measures (Fig. 1b,c), this pattern suggests long-term residency of eelgrass on the North American east coast and transport to the Northeast Atlantic via the North Atlantic Drift.

### Reticulated topology of *Z. marina* phylogeography

To further explore the degree of admixture and secondary contact, we constructed a split network^22^ using all ZM_Core_SNPs. Pacific populations were connected in a web-like fashion (Fig. 3a). WAS and BB were involved in alternative network edges (Fig. 3b), either clustering with SD or with both JS and JN. The topology places WAS and BB in an admixture zone with a northern Alaska component (ALI and ASL), and a more divergent southern component from SD, in line with the STRUCTURE results (Fig. 2c). In the Atlantic (Fig. 3c), edges among locations were shorter than those on the Pacific side, indicating a more recent divergence among Atlantic populations. A bifurcating topology connected the older Mediterranean populations, while both Northeast and Northwest Atlantic were connected by unresolved, web-like edges, indicating a mixture of incomplete lineage sorting and probable, recent gene flow.

**Fig. 3.**
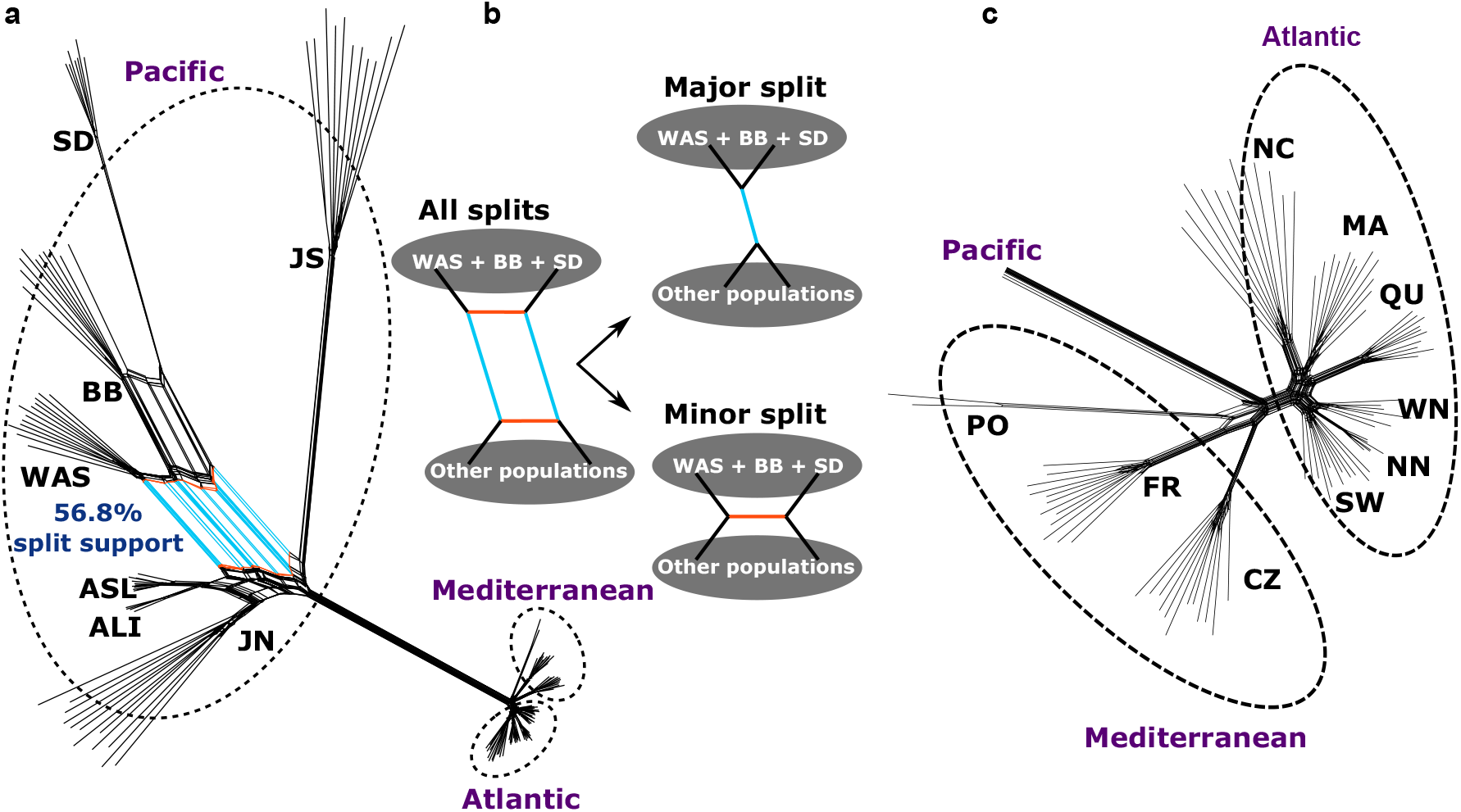
Conflicting phylogenetic signals in the nuclear genome. **a**, Splits network based on the core chromosomal SNP set (11,705 SNP, Supplementary Fig. 1). Each terminal branch indicates one individual sample. Splits colored in cyan are particularly strongly supported between a grouping of WAS, BB and SD and the rest of the Pacific. **b**, Main signals in the observed network structure. The splits network structure indicates that the SNP dataset supports alternative evolution histories, which are particularly strong with respect to BB, WAS and SD. The major split depicted in **b** is supported by 56.8% of all splits. **c**, Splits network reconstructed for Atlantic populations only.

We used Patterson’s D-statistic^23^ to further test for admixture^24^ (Supplementary Fig. 9). For the Pacific side, the pairs WAS/SD, BB/ALI, BB/ASL and to a lesser extent JN/ALI, showed the highest D-values (D=0.67; P<0.001), suggesting past admixture. For the Atlantic side, the pattern of admixture was less complex than in the Pacific, indicating recent or ongoing connection between Atlantic and Mediterranean Sea. This result is consistent with the admixture signal detected by STRUCTURE (SW, Fig. 2f), and with one Atlantic (SW) cpDNA haplotype that clusters with the Mediterranean ones (Fig. 2g).

### Time-calibrated multi-species coalescent (MSC) analysis and estimated times of major colonization events

Application of the multi-species coalescent^9^ (Fig. 4) assumes that populations diverge under a bifurcating model. Hence, locations that showed strong admixture (BB and WAS; Supplementary Fig. 9) were excluded from constructing a time-calibrated tree, leaving 14 populations. We further verified the dating of major events by additional exclusion of population involved in admixture (leaving seven populations), and found that time estimates for major divergence events were largely similar (Supplementary Fig. 10). Hence, we focused on the more comprehensive larger data set comprising 14 populations (Supplementary Fig.11).

**Fig. 4.**
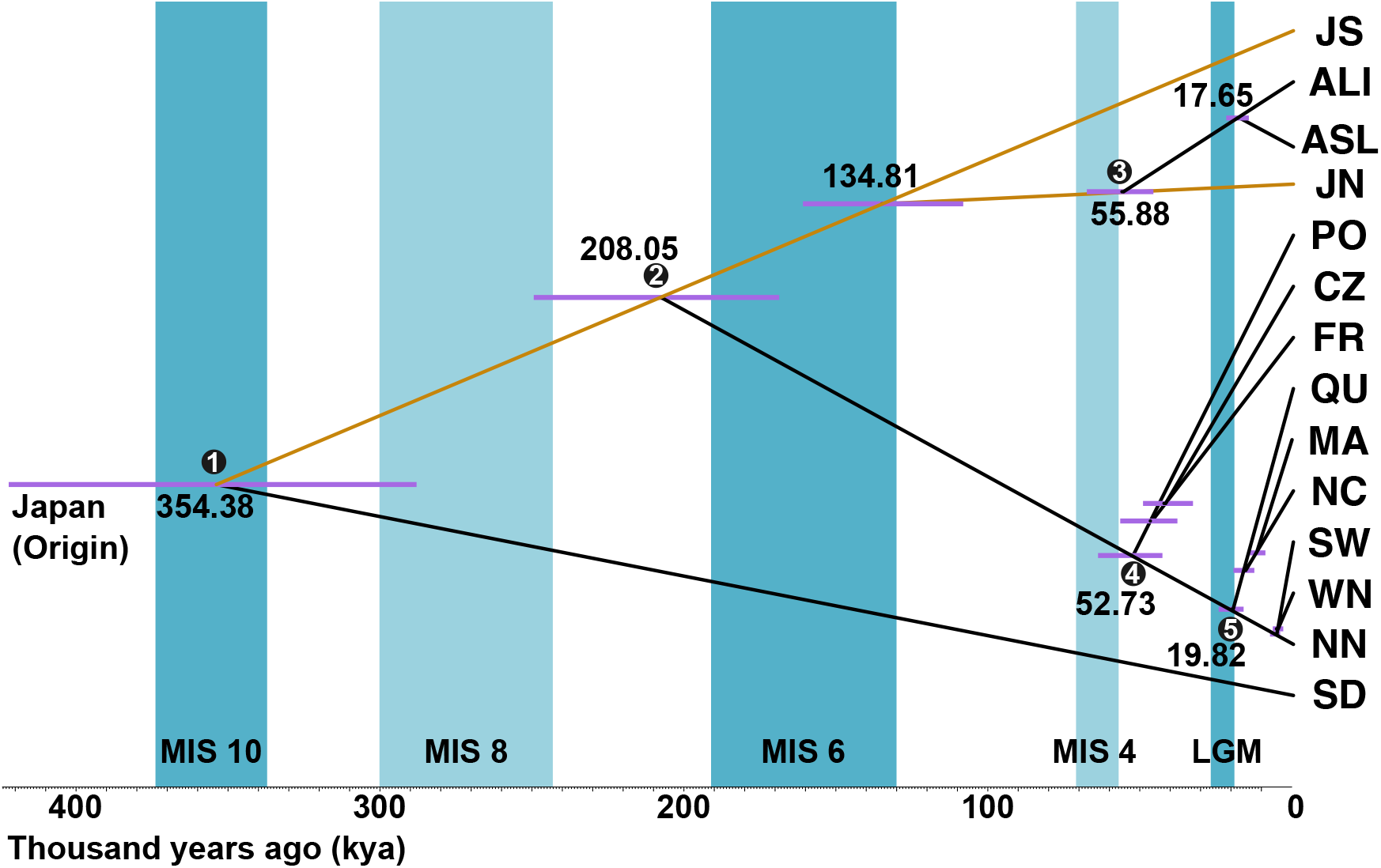
Time-calibrated phylogenetic tree based on the multi-species coalescent (MSC) allows dating of major colonization events. **a**, Blue bars indicate glacial periods with Marine Isotope Stages (MIS) alternating with warm to cool interglacial periods (white). Intensity of blue color depicts the intensity of glaciations. The Last Glacial Maximum (MIS2=LGM) is depicted at 26.5-19 kya. Estimated absolute divergence times of 7 nodes with stable topology (Supplementary Fig. 11) along with 95% confidence intervals (highest posterior densities, purple bars) are given. The two most strongly admixed populations WAS and BB were excluded (See Fig. 2 and 3). The orange edge connects the hypothetical founder in the Japan area with the extant JN and JS sites. Inferred colonization scenarios (numbered black dots on the nodes) are presented in Fig. 6.

As direct fossil evidence is unavailable within the genus *Zostera*, the divergence time between *Z. marina* and *Z. japonica* was estimated based on a calibration point that takes advantage of a whole-genome duplication event previously identified and dated to ∼67 mya^19^. The resulting clock rate for 4-fold degenerative transversions (4DTv) of paralogous gene sequences yielded a divergence time estimate of 9.86-12.67 mya between *Z. marina* and *Z. japonica* (Supplementary Note 2). We then repeated the analysis based on 13,732 SNP sites polymorphic within our target species (Supplementary Fig. 2) after setting a new *Z. marina*-specific calibration point.

Assuming JS as representative of the species origin^4^, we found direct evidence for two trans-Pacific dispersal events and indirect evidence for a third one (Fig. 4). The first trans-Pacific dispersal event at ∼354 Kya (95% highest posterior density HPD: 422-288 Kya) founded populations close to San Diego (SD) that remained isolated, but engaged in admixture to the north. Because dispersal from the West Pacific to the Atlantic requires stepping stones in the Northeast Pacific / Beringia, we infer a second trans-Pacific dispersal event from JN to the Northeast Pacific somewhat before *Z. marina* reached the Atlantic through the Canadian Arctic ∼209 Kya (95% HPD: 249-169 Kya). This estimate is surprisingly recent given that the Bering Strait opened as early as 4.8-5.5 mya ago^25^. The current Alaskan population (ASL) showed a strong signal of a recent 3rd trans-Pacific dispersal event from Japan that happened ∼55.9 Kya (95% HPD: 67.4-55.5 Kya), indicating (partial) replacement of *Z. marina* in Alaska with the new, extant populations. Further support comes from JN showing the smallest pairwise *F*_ST_ with all Atlantic populations (Supplementary Table 5). Moreover, JN was the only Pacific population that displayed a shared genetic component with the Atlantic (Fig. 2b).

In the Atlantic, divergence time estimates were much more recent than in the Pacific. The Mediterranean Sea clade emerged ∼52.7 Kya (95% HPD: 63.7-42.5 Kya). The Northwest and Northeast Atlantic also diverged very recently at ∼19.8 Kya (95% HPD: 24.1-15.8 Kya), and shared a common ancestor during the LGM, indicating that they were partially derived from the same glacial refugium in the Northwest Atlantic (likely at or near NC). Some admixture found in the Swedish (SW) population stemming from the Mediterranean gene pool (Fig. 2f,g) likely explains a higher genetic diversity at that location (Fig. 1b,c).

In a second coalescent approach^8^, we used alignments of 617 core genes across all samples (Supplementary Note 2). Based on the same initial calibration as under the multi-species coalescent, the tree topology was examined using ASTRAL while the time estimation was performed with StarBEAST2 (ref ^26^). This approach resulted in more recent divergence time estimates for the deeper nodes, while the more recent estimates were nearly identical (Supplementary Note 3, Supplementary Fig. 12,13).

Finally, we used the mutational steps among chloroplast (cpDNA) haplotypes as an alternative dating method. SD and BB along the Pacific East coast showed very different haplotypes, separated by about 30 mutations from the other Pacific and the Atlantic clades. Assuming a synonymous cpDNA mutation rate of 2*10^−9^ per site per year, this genetic distance corresponds to a divergence time of 392 Kya (Supplementary Note 4), comparable to the estimate of 354 Kya in the coalescent analysis. Conversely, few mutations (4-7) distinguished major Atlantic haplotypes from the Mediterranean Sea, consistent with recent divergence estimate based on nuclear genomes (Fig. 4).

### Demographic history and post LGM recolonization

We used the Multiple Sequentially Markovian Coalescent (MSMC)^27^ to infer past effective population size *N*_e_ (Fig. 5). Almost all eelgrass populations revealed a recent expansion 1,000-100 generations ago, while the magnitude of *N*_e_-value minima at about 10,000 to 1,000 generations varied. Given a range of plausible generation times of 3-5 yrs under a mix of clonal and sexual reproduction, is likely that the minimum *N*_e_ displayed by several locations coincides with the LGM. In general, low *N*_e_-values were related to a high degree of clonality at sites in northern (NN) and southern Europe (PO) (Supplementary Table 3). Within the Pacific Ocean, the southernmost population (SD), showed no drop in *N*_e_, while all others showed bottlenecks that became more pronounced from south to north (BB>WAS>ALI/ASL). As for the Atlantic side, the Northwest Atlantic populations NC/MA and the southern European populations PO/CZ (and to a lesser extent FR) showed little evidence for bottlenecks, suggesting that these localities represented refugia during the LGM. The opposite applied to QU in the Northwest and NN and SW in the Northeast Atlantic, where we see a pronounced minimal *N*_e_ at about 3,000-1,000 generations ago.

**Fig. 5.**
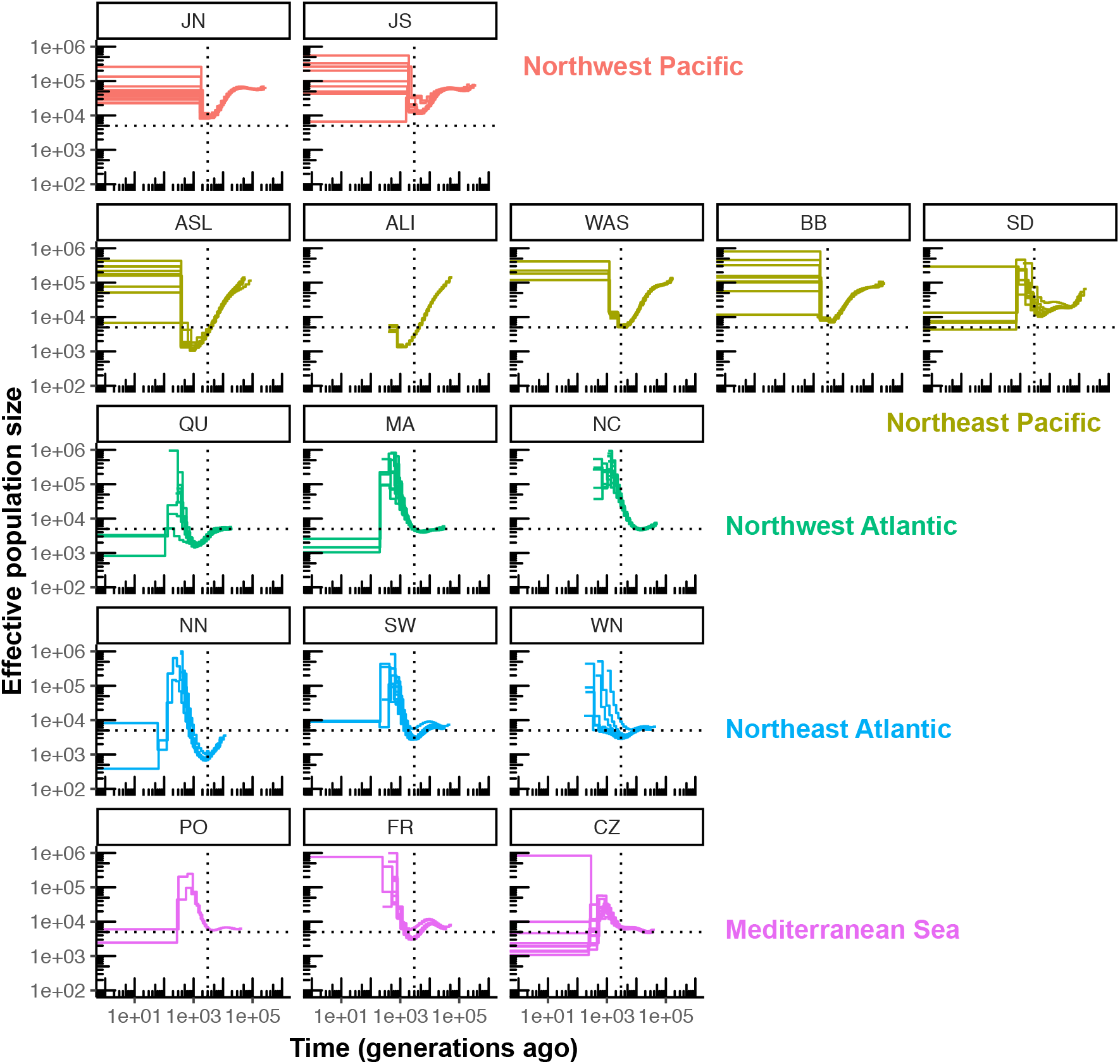
Demographic history of worldwide eelgrass (*Zostera marina*) populations reveal effects of the Last Glacial Maximum (LGM). Historical effective population sizes (*N*_e_) were inferred by the multiple sequentially Markovian coalescent (MSMC). Replicate runs were performed with all unique genotypes in each location, depicted as separate lines. The x-axis depicts generations rather than absolute time as generation time for *Z. marina* varies depending on the level of local clonality. *N*_e_-values are capped at 1 million. Many northern populations reveal a minimal *N*_e_ (thus likely a bottleneck) at ∼3,000 generations ago (dashed vertical lines), which probably reflects the impact of the LGM. Note that estimates younger than 1,000 generations are considered unreliable and are hence not be interpreted. The dashed horizontal lines at *N*_e_ = 5,000 are for orientation only.

For the Atlantic, we determined the most likely post-LGM recolonization through approximate Bayesian computations (ABC) (Supplementary Fig. 14) and found that areas around NC were the most likely glacial refugia for both the West and Northeast Atlantic locations.

## Discussion

In the current period of rapid climate change, the analysis of past climatic shifts and their legacy effects on genetic structure and diversity of extant populations is paramount^14,15,28^. *Z. marina* has a circumglobal distribution that provided us with the unique opportunity to reconstruct the natural expansion of a marine plant throughout the northern hemisphere starting from the species origin in the West Pacific during a period of strong recurrent climate changes (Fig. 6a,b).

**Fig. 6.**
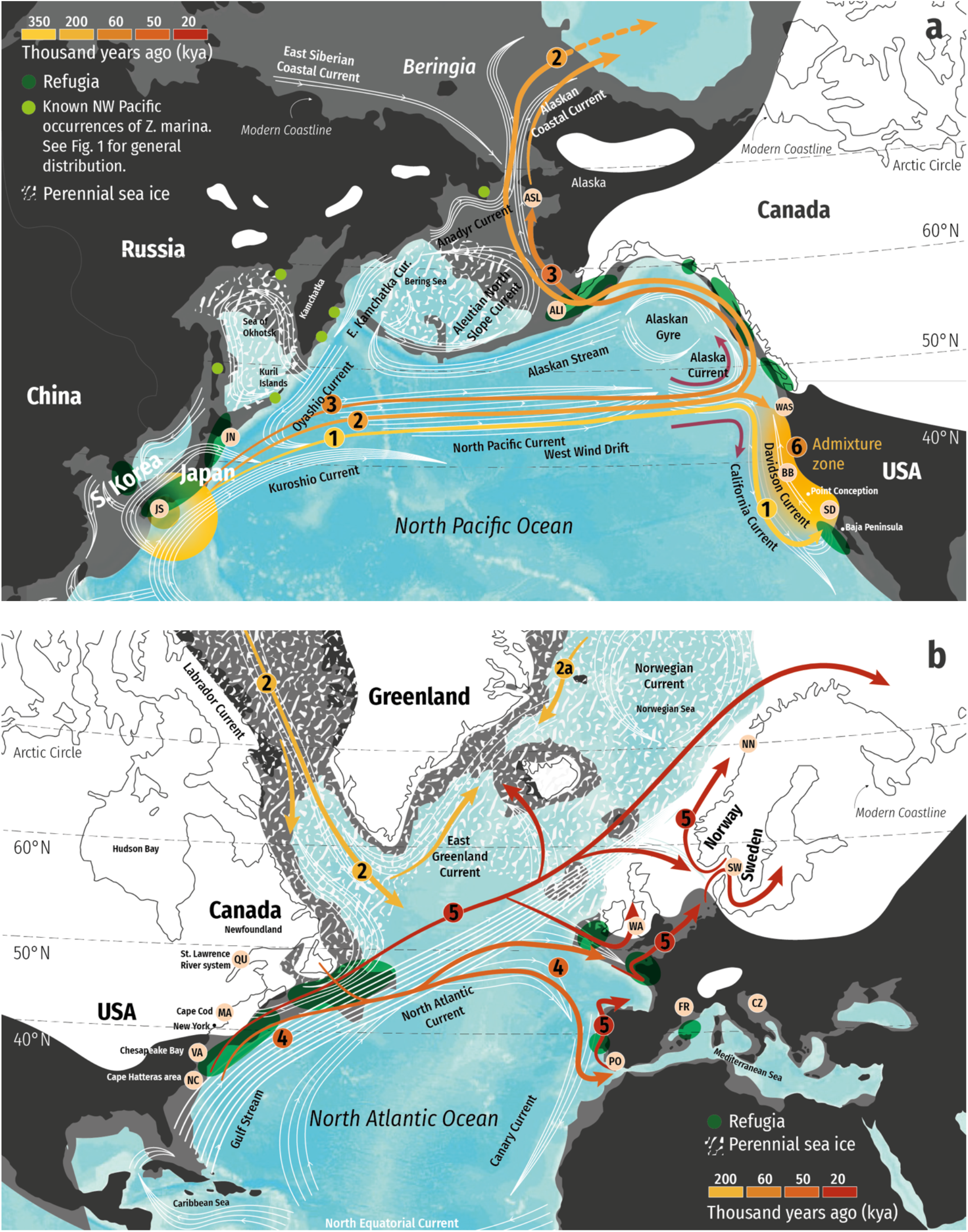
Dispersal and colonization history across the Pacific and to the Atlantic. For both maps: present coastline (black), LGM sea level coastline (dark gray), glaciers (white), perennial sea ice (speckled white), and current pathways (as shown). Sampled locations (pink dots with labels following Fig. 1), hypothesized refugia (dark green ovals). Dispersal pathways and timing (yellow-orange-red gradient arrows) including the North Pacific Current “gateway” (paired purple arrows). Numbers on current pathways correspond to phylogenetic branch points (nodes) in Fig. 4. **a**, Pacific Ocean. *Z. marina* arose in the Japanese Archipelago. Known occurrences in the Russian Arctic (light green dots). Hypothesized dispersal events: (1) first trans-Pacific dispersal via the North Pacific Current, arriving at the “gateway”, where it splits both south following the California Current, and north via the Alaska Current; (2) Second inferred trans-Pacific dispersal, ultimately arriving in the Atlantic, with an unknown, possibly extinct “ghost” population that was replaced by the extant Alaska population; (3) Alaska was colonized recently via North Japan in a third trans-Pacific event. SD ancestors may have later dispersed northwards (presumably via the Davidson Current), forming sequential admixtures with BB and WAS (“admixture zone”, event “6”). **b**, Atlantic Ocean. The dispersal into the Atlantic was likely propelled by the southward Labrador current (2). (4) original foundation of the Mediterranean populations (including Portugal) and further along the Atlantic coastlines with (5) post-LGM recolonization of the East Atlantic via refugia close to NC (and hypothesized southern European refugia), subsequent expansion northward as the ice retreated and shorelines formed.

The presence of eelgrass in the Atlantic is surprisingly recent, dating to only ∼208 Kya (95% HPD: 249-169 Kya). As no other seagrass species is able to fill this ecological niche or form dense meadows in boreal to Arctic regions (>50 °N, Supplementary Note 1), historical contingency^6^ has played a previously underappreciated role for the establishment of this unique and productive ecosystem. The recency of the arrival of eelgrass on both sides of the Atlantic may also explain why relatively few animals are endemic to eelgrass beds nor have evolved to consume its plant tissue directly, while most of the biomass produced ends up either in the sediment as blue carbon, or is exported into the detritus based food chain^29^. The first dated population-level phylogeny in any seagrass species might also explain why there seems to be little niche differentiation among eelgrass-associated epifauna in the Atlantic compared to the Pacific^30^. Our study demonstrates how macro-ecology, here the presence of an entire ecosystem, may be strongly determined by the colonization history, specifically the timeframe in which eelgrass reached the North Atlantic^6^, and not by suitable environmental conditions.

We identified the North Pacific Current that began to intensify ∼one million years ago^31^ as major dispersal gateway. San Diego (SD) was colonized by the earliest detectable colonization event roughly 400 Kya (Fig. 6a, event “1”), and has retained old genetic variation since then, probably owing to rarity of genetic exchange southward across the Point Conception biogeographic boundary^32^ and a weak and variable Davidson Current. Subsequent trans-Pacific events eventually resulted in an admixture zone in intermediate WAS and BB situated among the ancient SD clade and the younger Alaskan ones (ASL/ASI, Fig. 6a, event “6”).

The second trans-Pacific dispersal (Fig. 6a, event “2”) actually paved the way for an inter-oceanic dispersal, the colonization of the Atlantic through the Arctic Ocean, possibly via the stepping stone of an Arctic “ghost” population. The latter was replaced with more recent immigrant genotypes from northern Japan in a third detected dispersal from West to East Pacific (Fig. 6a, event “3”). Although the Bering Strait may have opened as early as 5.5-4.8 mya^25^, we were only able to detect a single colonization event into the Atlantic, in contrast to other amphi-Arctic and boreal marine invertebrates^33^ and seaweeds^34^. Genomic variation characteristic of extant Alaskan populations was not detected in any North-Atlantic populations, in line with earlier microsatellite data^35^, suggesting that the Atlantic was only colonized once. While we cannot rule out an earlier colonization, this would require that they became extinct without leaving any trace extant in nuclear genomes or cpDNA haplotypes, which we consider unlikely.

The Pacific-Atlantic genetic divide was recently identified as a “Pleistocene legacy” based on a marker-based genotyping study^15^. Here, we demonstrate the presence of two deeply divergent clades in the Pacific that share a complex pattern of secondary contact on the East Pacific side (Supplementary Note 5). In contrast, a clear genetic separation between West and East Atlantic populations is not evident suggesting recent population contractions and expansions driven by the LGM, with the North Atlantic Drift driving repeated west-east colonization events (Fig. 6b).

While our phylogeny (Fig. 4) would also be consistent with a scenario in which the deep branching SD population would represent the origin of *Z. marina*, we consider this very unlikely given the prevailing ocean currents (Fig. 6a), the patterns of genetic diversity (Fig. 1b,c) and our current understanding of the emergence of the genus *Zostera* (∼15 mya), including the species *Z. marina* some 5-1.62 mya^4^in the Northwest Pacific. Other *Zostera* species have also been seeded to other parts of the globe by multiple dispersal events from the genus-origin close to Japan^4^. Thus, considering all evidence jointly, we conclude that Japan, and not the East Pacific (SD), is the most likely geographic origin of eelgrass and the source of multiple dispersal events with ocean currents.

Two major LGM refugia were detected in the Atlantic, of which one near North Carolina (NC) apparently served as source population for the entire Northwest and Northeast Atlantic (Fig. 6b, event “5”), as in other marine species^11,36^ including seaweeds^37^. Additionally, the Mediterranean Sea was a refugium itself. We may have missed a role of Brittany to be a refugium, as has been reported for seaweeds and invertebrates^37,38^, as it was not sampled.

Along with demographic modeling we identify population contraction and subsequent latitudinal expansion along three coastlines following the LGM (26-19 Kya). These are common patterns of many terrestrial^10^ and intertidal species^13,39^, with the Northeast Atlantic/North Sea coastline and Beringia being most drastically affected. Interestingly, for *Z. marina*, the Atlantic region was not more severely influenced by the last glaciations and sea level changes than the East Pacific (Fig. 5; 6b), relative to its much lower baseline diversity (Supplementary Table 4), while we are lacking the sample location to examine this for the West Pacific. This ultimately resulted in dramatic differences in genome-wide diversity. The 5-to 7-fold lower overall genetic diversity in the Atlantic adds to marked LGM effects and resulted in >30-fold differences among populations with the highest (JS) vs. lowest (NN) diversity, with currently unknown consequences for the adaptive potential and genetic rescue of eelgrass in the anthropocene.

The relatively low number of extant seagrass species (ca. 65 species in six families^40^) has been attributed to frequent intermediate extinctions^5^. Our data suggest a second plausible process, namely multiple long-distance genetic exchange among ocean basins that may have impeded allopatric speciation (see also^41^). Our range-wide sampling has allowed an overview of evolutionary history in this lineage of seagrass and opens the door for exploration of functional studies across ocean basins and coasts. Future work will explore the pan-genome of *Z. marina* with the consideration of how the high diversity and robustness of Pacific populations may contribute to management and rescue of populations along rapidly warming Atlantic coastlines.

## Online Methods

### Study species and sampling design

Our study species eelgrass (*Zostera marina* L.) is the most widespread seagrass species of the temperate to Arctic northern hemisphere. It is being developed as model for studying seagrass evolution and genomics (e.g.;^15,17,19,42^. *Z. marina* is a foundation species of shallow water ecosystems^15^ with a number of critical ecological functions including enhancing the recruitment of fish and crustaceans^43^, improvement of water quality^44^ and the sequestration of “blue carbon”^45^.

Eelgrass features a mix of clonal (=vegetative) and sexual reproduction, with varying proportions across sites^39^. Hence, in most populations, except for the most extreme cases of mono-clonality^46^, replicated modular units (leaf shoots= ramets) stemming from a sexually produced individual (=genet or clone) are intermingled to form the seagrass meadow. This also implies that generation times are difficult to estimate or average across populations.

We conducted a range-wide sampling collection of 190 *Z. marina* specimen from 16 geographic populations (Fig. 1a; Supplementary Table 1). The chosen locations were a subset of the *Zostera* Experimental Network (ZEN) sites that were previously analyzed using microsatellite markers^15^. Although a sampling distance of >2 m was maintained to reduce the likelihood of collecting the same clone twice this was not always successful (cf. Supplementary Table 3 which also provides estimates of local clonal diversity). Plant tissue was selected from the basal meristematic part of the shoot after peeling away the leaf sheath to minimize epiphytes (bacteria and diatoms), frozen in liquid nitrogen and stored at -80 °C until DNA extraction.

### DNA extraction, whole-genome resequencing and quality check

Genomic DNA was extracted using the Macherey-Nagel NucleoSpin plant II kit following the manufacturer’s instructions. Hundred-200 mg fresh weight of basal leaf tissue, containing the meristematic region was ground in liquid N_2_. DNA concentrations were in the range of 50-200 ng/uL. Quality control was performed following JGI guidelines (https://jgi.doe.gov/wp-content/uploads/2013/11/Genomic-DNA-Sample-QC.pdf). Plate-based DNA library preparation for Illumina sequencing was performed on the PerkinElmer Sciclone NGS robotic liquid handling system using Kapa Biosystems library preparation kit. Two hundred ng of sample DNA were sheared to a length of around 600 bp using a Covaris LE220 focused-ultrasonicator. Selected fragments were end-repaired, A-tailed, and ligated with sequencing adaptors containing a unique molecular index barcode. Libraries were quantified using KAPA Biosystems’ next-generation sequencing library qPCR-kit on a Roche LightCycler 480 real-time PCR instrument. Quantified libraries were then pooled together and prepared for sequencing on the Illumina HiSeq2500 sequencer using TruSeq SBS sequencing kits (v4) following a 2x150 bp indexed run recipe to a targeted depth of approximately 40x coverage. The quality of the raw reads was assessed by FastQC (https://www.bioinformatics.babraham.ac.uk/projects/fastqc/) and visualized by MultiQC^47^. BBDuk (https://jgi.doe.gov/data-and-tools/bbtools/bb-tools-user-guide/bbduk-guide/) was used to remove adapters and for quality filtering, discarding sequence reads (i) with more than one “N” (maxns=1); (ii) shorter than 50 bp after trimming (minlength=50); (iii) with average quality <10 after trimming (maq=10). FastQC and MultiQC were used for second round of quality check for the clean reads. Sequencing coverage was calculated for each sample (Supplementary Data 1).

### Identifying core and variable genes

In order to analyze genetic loci present throughout the global distribution range of eelgrass, we focused on identifying core genes that would be present in genomes of all individuals. To do so, each of the 190 ramets were *de novo* assembled using HipMer (k=51)^48^. To categorize, extract, and compare core and variable (shell and cloud) genes, primary transcript sequences (21,483 gene models) from the *Z. marina* reference (V3.1) ref^17^ were aligned using BLAT using default parameters^49^ to each *de novo* assembly. Genes were considered present if the transcript aligned with either (i) >60% identity and >60% coverage from a single alignment, or (ii) >85% identity and > 85% coverage split across three or fewer scaffolds. Individual presence-absence-variation (PAV) calls were combined into a matrix to classify genes into core, cloud, and shell categories based on their observation across the population. The total number of genes considered was 20,100. Because identical genotypes and fragmented, low-quality assemblies can bias and skew PAV analyses, only 141 single representatives of clones and ramets with greater than 17,500 genes were kept to ensure that only unique and high-quality assemblies were retained. Genes were classified using discriminant analysis of principal components (DAPC)^50^ into cloud, shell, and core gene clusters based on their frequency. Core genes were the largest category, with 18,717 genes that were on average observed in 97% of ramets.

### SNP mapping, calling and filtering

The quality-filtered reads were mapped against the chromosome-level *Z. marina* reference genome V3.1 using BWA MEM^51^. The alignments were converted to BAM format and sorted using Samtools^51^. The MarkDuplicates module in GATK4^52^ was used to identify and tag duplicate reads in the BAM files. Mapping rate for each genotype was calculated using Samtools (Supplementary Data 2). HaplotypeCaller (GATK4) was used to generate a GVCF format file for each sample, and all the GVCF files were combined by CombineGVCFs (GATK4). GenotypeGVCFs (GATK4) was used to call genetic variants.

BCFtools^53^ was used to remove SNPs within 20 base pairs of an indel or other variant type (Supplementary Fig. 1) as these variant types may cause erroneous SNPs calls. VariantsToTable (GATK4) was used to extract INFO annotations. SNPs meeting one or more than one of the following criteria were marked by VariantFiltration (GATK4): MQ < 40.0; FS > 60.0; QD < 10.0; MQRandSum > 2.5 or MQRandSum < -2.5; ReadPosRandSum < -2.5; ReadPosRandSum > 2.5; SOR > 3.0; DP > 10804.0 (2 * average DP). Those SNPs were excluded by SelectVariants (GATK4). A total of 3,975,407 SNPs were retained. VCFtools^54^ was used to convert individual genotypes to missing data when GQ <30 or DP <10.

Individual homozygous reference calls with one or more than one reads supporting the variant allele, and individual homozygous variant calls with ≥ 1 read supporting the reference were set as missing data. Only bi-allelic SNPs were kept (3,892,668 SNPs). To avoid the reference-related biases owing to the Pacific-Atlantic genomic divergence, we focused on the 18,717 core genes that were on average observed in 97% of ramets. Bedtools^55^ was used to find overlap between the SNPs and the core genes, and only those SNPs were kept (ZM_HQ_SNPs, 763,580 SNPs). Genotypes that were outside our custom quality criteria were represented as missing data.

### Excluding duplicate genotypes, genotypes originating from selfing, and those with high missing rate

Based on the extended data set ZM_HQ_SNPs (763,580 SNPs; Supplementary Fig. 1), possible parent-descendant pairs under selfing (Supplementary Table 2) as well as clonemates were detected based on the shared heterozygosity (SH)(ref^56^). To ensure that all genotypes assessed originated by random mating, ten ramets showing evidence for selfing were excluded. Seventeen multiple sampled clonemates were also excluded (Supplementary Table 3, Supplementary Fig. 3). Based on ZM_HQ_SNPs (763,580 SNPs), we calculated the sample-wise missing rate using a custom Python3 script and plotted results as a histogram (Supplementary Fig. 4). Missing rates were mostly <15%, except for ten ramets (ALI01, ALI02, ALI03, ALI04, ALI05, ALI06, ALI10, ALI16, QU03, and SD08) that were also excluded.

### Chloroplast haplotypes

For the chloroplast analysis, 28 samples were excluded owing to evidence for selfing and membership to the same clone, while lack of coverage was not an issue. Chloroplast genome was de novo assembled by NOVOPlasty^57^. The chloroplast genome of *Z. marina* was represented by a circular molecule of 143,968 bp with a classic quadripartite structure: two identical inverted repeats (IRa and IRb) of 24,127 bp each, large single-copy region (LSC) of 83,312 bp, and small single-copy region (SSC) of 12,402 bp. All regions were equally taken into SNP calling analysis except for 9,818 bp encoding 23S and 16S RNAs due to supposed bacteria contamination in some samples. The raw Illumina reads of each individual were aligned by BWA MEM to the assembled chloroplast genome. The alignments were converted to BAM format and then sorted using Samtools^51^. Genomic sites were called as variable positions when frequency of variant reads >50% (Supplementary Fig. 8) and the total coverage of the position >30% of the median coverage (174 variable positions). Then 11 positions likely related to microsatellites and 12 positions reflecting minute inversions caused by hairpin structures^58^ were removed from the final set of variable positions for the haplotype reconstruction (151 SNPs).

### Putatively neutral and non-linked SNPs

Among a total of 153 unique samples that were retained for analyses, SnpEff (http://pcingola.github.io/SnpEff/) was used to annotate each SNP. To obtain putatively neutral SNPs, we kept only SNPs annotated as “synonymous_variant” (ZM_Neutral_SNPs, 144,773 SNPs). For the SNPs in ZM_Neutral_SNPs (144,773 SNPs), only SNPs without any missing data were kept. To obtain putatively non-linked SNPs, we thinned sites using Vcftools to achieve a minimum pairwise distance (physical distance in the reference genome) of 3,000 bp to obtain our core data set, hereafter ZM_Core_SNPs, corresponding to 11,705 SNPs.

### Genetic population structure based on nuclear and chloroplast polymorphism

We used R-packages to run a global principal component (PCA) analysis based on ZM_HQ_SNPs, (=763,580 SNPs). The package vcfR^59^ was used to load the VCF format file, and function glPca in adegenet package to conduct PCA analyses, followed by visualization through the ggplot2 package. We used Bayesian clustering implemented in STRUCTURE to study population structure and potential admixture^20^. To reduce the run time, we randomly selected 2,353 SNPs from ZM_Core_SNPs (20%) to run STRUCTURE (Length of burn-in period 3*10^5^; number of MCMC runs 2*10^6^). Ten runs were performed for K-values 1-10. StructureSelector^60^ was used to decide the optimal K based on Delta-K method^21^, and to combine and visualize the STRUCTURE results of 10 independent runs for each K-value in this and the subsequent analyses.

In order to detect nested population structure, the global run was complemented with analyses of populations from the Atlantic and Pacific side, respectively. Pacific data were extracted from ZM_Neutral_SNPs (144,773 SNPs), excluding monomorphic sites and those with missing data. To obtain putatively independent SNPs, we thinned sites using Vcftools, so that no two sites were within 3,000 bp distance (physical distance in the reference genome) from one another (ZM_Pacific_SNPs, 12,514 SNPs). Those 12,514 SNPs were used in the PCA, while a set of randomly selected 6,168 SNPs was used in STRUCTURE to reduce run times (Length of burn-in period 3*10^5^; number of MCMC runs 2*10^6^) as described above and with possible K-values between 1 and 7.

Polymorphism data for Atlantic and Mediterranean eelgrass were extracted from ZM_Neutral_SNPs (144,773 SNPs). To obtain putatively independent SNPs, we thinned sites using Vcftools according to the above criteria. The resulting 8,552 SNPs were then used to run another separate PCA and STRUCTURE using the parameters above. For STRUCTURE analysis, K was set from 1 to 5. For each K, we repeated 10 times independently (Supplementary Fig. 6,7).

For the cpDNA data, the population structure was explored using a haplotype network, constructed via the Median Joining Network method^61^ with epsilon 0 and 1 implemented by PopART^62^, based on 151 polymorphic sites.

### Analysis of reticulate evolution using split network

To assess reticulate evolutionary processes, we used SplitsTree4^22^ to construct a split network, which is a combinatorial generalization of phylogenetic trees and is designed to represent incompatibilities. A custom Python3 script was used to generate a fasta format file containing concatenated DNA sequences for all ramets based on ZM_Core_SNPs. For a heterozygous genotype, one allele was randomly selected to represent the site. The fasta format file was converted to nexus format file using MEGAX^63^, which was fed to SplitsTree4. NeighborNet method was used to construct the split network.

### Genetic diversity

Vcftools was used to calculate nucleotide diversity (π) for each population at all synonymous sites using each of the six chromosomes as replicates for 44,685 SNPs without any missing data (Supplementary Fig. 1). Genomic heterozygosity for a given genotype *H*_OBS_ (as number of heterozygous sites) / (total number of sites with available genotype calls) was calculated using a custom Python3 script based on all synonymous SNPs (144,773).

### Pairwise population differentiation using *F*_ST_

We used the function stamppFst in the StAMPP-R package^64^ to calculate pairwise *F*_ST_ based on ZM_Core_SNPs (Supplementary Table 4). *P*-values were generated by 1,000 bootstraps across loci.

### D-statistics

Patterson’s D provides a simple and powerful test for the deviation from a strict bifurcating evolutionary history. The test is applied to three populations P1, P2, and P3 plus an outgroup O, with P1 and P2 being sister populations. If P3 shares more derived alleles with P2 than with P1, Patterson’s D will be positive. We used Dsuite^24^ to calculate D-values for populations within the Pacific and Atlantic side, respectively (Supplementary Fig. 9), respectively. D was calculated for trios of *Z. marina* populations based on the SNP core dataset (ZMZJ_D_SNPs) (Supplementary Fig. 2), using *Z. japonica* as outgroup. The Ruby script plot_d.rb (https://github.com/mmatschiner/tutorials/blob/master/analysis_of_introgression_with_snp_data/src/plot_d.rb) was used to plot a heatmap that jointly visualizes both the D-value and the associated p value for each comparison of P2 and P3. The color of the corresponding heatmap cell indicates the most significant D value across all possible populations in position P1. Red colors indicate higher D values, and more saturated colors indicate greater significance.

### Phylogenetic tree with estimated divergence time

To estimate the divergence time among major groups, we used the multi-species coalescent in combination with a strict molecular clock model^9^. We used the software SNAPP^7^ with an input file prepared by script “snapp_prep.rb” (https://github.com/mmatschiner/snapp_prep). Two specimen were randomly selected from each of the included populations, and genotype information was extracted from ZMZJ_Neutral_SNPs (Supplementary Fig. 1,2).

Monomorphic sites were excluded. Only SNPs without any missing data were kept. To obtain putatively independent SNPs, we thinned sites using Vcftools, so that no two included SNPs were within 3,000 bp (physical distance in the reference genome) from one another (6,169 SNPs). The estimated divergence time between *Z. japonica* and *Z. marina* was used as calibration point, which was implemented as a lognormal prior distribution (Supplementary Note 2, mean = 11.154 mya, SD = 0.07).

A large proportion of the 6,169 SNPs above represented the genetic differences between *Z. japonica* and *Z. marina*, and were monomorphic in *Z. marina*. To obtain a better estimation among *Z. marina* populations, we performed a second, *Z. marina*-specific SNAPP analysis via subsampling from the ZM_Neutral_SNPs (144,773 SNPs) data set, excluding monomorphic sites and missing data. We thinned sites again using Vcftools, so that all sites were ≥3,000 bp distance from one another (13,732 SNPs). The crown divergence for all *Z. marina* populations, estimated in the first SNAPP analysis, was used as calibration point, and implemented as a lognormal prior distribution (mean = 0.3564 Mya, SD = 0.1).

As the multi-species coalescent model does not account for genetic exchange, the SNAPP analysis was repeated after removing certain populations based on admixture assessments via STRUCTURE (Fig. 2), SplitsTree (Fig. 3) and D statistics (Supplementary Fig. 9). This produced two reduced data sets: The first included seven populations from which for the Pacific side, WAS, BB, and ALI were excluded, while for the Atlantic side, NC, SW, and CZ were selected to be representatives for the Northwest Atlantic, Northeast Atlantic, and the Mediterranean Sea, respectively (Supplementary Fig. 11). Here, we focus on a more complete set with 14 populations where only two Pacific locations WAS and BB (involved in admixture with SD) were excluded. This was legitimate as time estimates for major divergence events were very similar (compare Fig. 4. to Supplementary Fig. 11).

### Demographic analysis

The Multiple Sequentially Markovian Coalescent^27^ was run for each genotype per population. We here focus on time intervals where different replicate runs per population converged, acknowledging that MSMC creates unreliable estimates in recent time^65^. Owing to marked differences in the degree of clonality and the relative amount of sexual vs. clonal reproduction, the generation time of *Z. marina* varies across populations which prevented us to represent the x-axis in absolute time.

We first generated one mappability mask file for each of the six main chromosomes using SNPable (http://lh3lh3.users.sourceforge.net/snpable.shtml). Each file contained all regions on the chromosome that permitted unique mapping of sequencing reads. We then generated one mask file for all core genes along each of the six main chromosomes. We generated one ramet-specific mask file based on the bam format file using bamCaller.py (https://github.com/stschiff/msmc-tools), containing the chromosomal regions with sufficient coverage of any genoytpe. The minDepth variable in bamCaller.py was set to 10. We also generated a ramet-specific vcf file for each of the six main chromosomes based on ZM_HQ_SNPs using a custom Python3 script.

### Recolonization scenarios after the LGM for the Atlantic

DIYABC-RF^66^ was used to run simulations under each scenario to distinguish between alternative models of the recolonization history of *Z. marina* after the LGM. Considering that the Mediterranean Sea had its own glacial refugium, the ABC-modeling was conducted for only the Atlantic. We constructed three recolonization scenarios (Supplementary Fig. 12) (i) NC and MA were glacial refugia in the Atlantic, first recolonized QU as stepping stone and then the Northeast Atlantic. (ii) NC and MA represent the only glacial refugia in the Atlantic. Both QU and Northeast Atlantic were directly recolonized by the glacial refugia. (iii) NC and MA represent the southern glacial refugia for the Northwest Atlantic only.

## Supporting information

Sequence coverage

Sample mapping rate

Short read archive accession numbers

## Data and code availability

Genome data have been deposited in Genbank (short read archive, Supplementary data 3). Coding sequences of *Z. japonica* and *Z. marina* for the ASTRAL analysis can be found on figshare (doi.org/10.6084/m9.figshare.21626327.v1). VCF files of the 11,705 core SNPs can be accessed at doi.org/10.6084/m9.figshare.21629471.v1. Custom-made scripts were deposited on GitHub (github.com/leiyu37/populationGenomics_ZM.git).

## Acknowledgements

This study was supported by a PhD-scholarship from the China Scholarship Council (CSC) to L.Y. (No. 201704910807), by a fellowship to M.K in the Helmholtz School for Marine Data Science (MarDATA, grant no HIDSS-0005), and by a grant to Jonathan Eisen, J.J.S. and J.L.O from the U.S. Department of Energy (DOE) Joint Genome Institute (JGI) Community Sequencing Program (CSP 502951, 2016, Population and evolutionary genomics of host-microbiome interactions in *Zostera marina* and other seagrasses). The work (proposal: 10.46936/10.25585/60000773) conducted by the U.S. Department of Energy Joint Genome Institute (https://ror.org/04xm1d337), a DOE Office of Science User Facility, is supported by the Office of Science of the U.S. Department of Energy operated under Contract No. DE-AC02-05CH11231. Field sampling was supported by the National Science Foundation (OCE-1336206 to JED). We thank X. Zhang for providing the unpublished reference genome of *Zostera japonica* to predict the coding sequences, Susanne Landis (scienstration) for assisting with figures and illustrations and the many other members of the *Zostera* Experimental Network (ZEN). We thank T. Bayer for discussions on bioinformatic problems and Y. Li for assistance with the ABC-RF analysis.

## Sampling permits and compliance with the Convention on Biological Diversity

All samples were obtained in compliance with national regulations for the sampling of biological material, including the adherence to the regulations laid out in the national guidelines to assure fair share of genomic information (“Nagoya”-protocol).

