## Supplementary_Notes_Tables_Figures for "Ocean currents drive the worldwide colonization of the most widespread marine plant, eelgrass (*Zostera marina*)"

### Supplementary Note 1 - Ecological niches of seagrass species

#### Ecological niches of temperate-to-Arctic North Atlantic seagrass species

Here we briefly review the ecological niches of Atlantic seagrass species in the temperate to Arctic northern hemisphere in which *Zostera marina* dominates the thermal range, spanning 18°C in average annual temperatures and 40° latitude (up to 72°N). Although mixed beds involving (usually two) seagrass species occur, *Z. marina* dominates and is typically monospecific, thus creating a unique foundational niche for which there is no substitute.

On the Northwestern Atlantic coast, the southern distribution limit of eelgrass in North Carolina, where it sometimes co-occurs with the subtropical species *Halodule wrightii*. The latter species reaches its northern distribution limit in North Carolina (Ferguson et al. 1993) and can co-occur with *Z. marina. Ruppia maritima* also occurs in mixed beds with *Z. marina* in parts of inner Chesapeake Bay (Koch & Orth 2003). All three species are exclusively found in lagoon setting and not along exposed shores. They colonize a wide range of salinities (from 10-20 psu to fully marine at 30-35 psu).

On the Northeastern Atlantic coast, eelgrass occurs as far south as southern Portugal where it sometimes intermingles with the more tropical *Cymodosa nodosa. C. nodosa* occurs only as far north as southern Portugal (42°N, Chefaoui et al. 2016) but is also widely distributed throughout the Mediterranean, sometimes mixed with patches of *Z. marina*. It cannot survive in winter temperatures <6°C. *Posidonia oceanica* is the iconic seagrass species endemic to the Mediterranean (Procaccini et al. 2001). It rarely forms mixed beds with *Z. marina.*

*Nanozostera noltii* (also known as *Zostera noltii* in the ecological literature) is a Tethyan species arriving in the Mediterranean and spreading to inland seas of the region some 14-11 mya. It survived the Mediterranean salinity crisis (6-5 mya). It occurs along East Atlantic shores as far as Northern Scotland and southern Norway (56°N). Although it occurs in mixed stands with *Z. marina* in some areas (Laugier et al. 1999), its small size (maximal canopy height 15 cm) generally confines it to the intertidal zone and cannot provide the same habitat function as dense and large lower intertidal and subtidal meadows of *Zostera marina*. When occurring together, *Z. marina* is the canopy forming species attaining about 3-times the plant height compared to *Nanozostera noltii*.

Chefaoui RM, Assis J, Duarte CM, Serrão EA (2016) Large-Scale Prediction of Seagrass Distribution Integrating Landscape Metrics and Environmental Factors: The Case of *Cymodocea nodosa* (Mediterranean–Atlantic). Estuaries and Coasts 39: 123-137 doi 10.1007/s12237-015-9966-y

Ferguson RL, Pawlak BT, Wood LL (1993) Flowering of the seagrass *Halodule wrightii* in North Carolina, USA. Aquatic Botany 46: 91-98 doi https://doi.org/10.1016/0304-3770(93)90066-6

Koch, EW, Orth RJ. (2003) The Seagrasses of the Mid-Atlantic coast of the United States, *in* Green EP, Short FT. (2003) World Atlas of Seagrasses. Univ. California Press.

Laugier T, Rigollet V, de Casabianca M-L (1999) Seasonal dynamics in mixed eelgrass beds, *Zostera marina* L. and *Z. noltii* Hornem., in a Mediterranean coastal lagoon (Thau lagoon, France). Aquatic Botany 63: 51-69 doi https://doi.org/10.1016/S0304-3770(98)00105-3

Procaccini G, Orsini L, Ruggiero MV, Scardi M (2001) Spatial patterns of genetic diversity in *Posidonia oceanica*, an endemic Mediterranean seagrass. Mol Ecol 10: 1413-1421

#### Ecological niches of the temperate-to-Arctic North Pacific seagrass species

On the NE temperate-to-Arctic Pacific coast, *Z. marina* dominates soft bottoms in monospecific stands from Northern Alaska to Baja California Mexico (Wyllie-Echeverria and Ackerman 2003). *Ruppia maritima* has the same distribution but is more restricted to lower salinity lagoons where the species can form mixed beds with *Z. marina*. *Zostera japonica* and *Zostera asiatica* are human introductions from the early 20$^{th}$ century. Three species of *Phyllospadix* (known as surfgrasses) are found from mid-Alaska to Baja but dominate the rocky surf zone as their unique niche. The subtropical *Halodule wrightii* is occasionally encountered and restricted to the Gulf of California, Mexico

The NW temperate-to-Arctic Pacific coast of mid-Japan northward is a diversity hotspot for seagrasses (Aioi and Nakaoka 2003) in general and *Zostera* in particular—with four species of *Zostera* and two of *Phyllospadix. Z. japonica* is a small plant found from SW Japan to Sakhalin and the East coast of Kamchatka, Russia. Mixed meadows of 3-4 seagrass species are common in both the tropical and temperate regions where zonation by depth plays a role; *Z, marina* is typically shallow subtidal in this assemblage.

Aioi K, Nakaoka M (2003). The seagrasses of Japan, *in* Green EP, Short FT. (2003) World Atlas of Seagrasses. Univ. California Press.

Wyllie-Echeverria S, Ackerman JD (2003) The Pacific coast of North America *in* Green EP, Short FT. (2003) World Atlas of Seagrasses. Univ. California Press.

### Supplementary Note 2 - rate estimation of the synonymous molecular clock in *Zostera marina*

#### Estimating the divergence time between *Zostera japonica* and *Z. marina*

To calibrate a molecular clock for *Z. marina*, we first calculated the fourfold degenerative third-codon transversion rate (4DTv) for each of the 1,072 syntenic paralog pairs in the *Z. marina* reference V3.1 (as determined with the comparative genomics pipeline GENESPACE; Schmutz & Lovell, 2021). The 4DTv rates showed a distinct peak at approximately 0.48, indicating the *Z. marina* whole genome duplication event (~67 mya; Olsen et al., 2016). The 90% confidence interval of the peak (0.4640-0.5164) was estimated based on 10,000 bootstraps.

The divergence time between *Z. marina* and *Z. japonica* was then estimated based on the divergence of homologs. A constrained protein homology search was performed using BLAT and GeMoMa (v1.7; Keilwagen et al., 2019). To generate a constrained sequence database to search, *Z. marina* transcripts were aligned to the *Z. japonica* genome assembly. Each transcript's best hit location (+500 bp sequence buffer) was extracted from the *Z. japonica* genome and was used for GeMoMa protein prediction with default parameters. The protein prediction pipeline found 8,687 peptide sequences. Gffread (Pertea & Pertea, 2020) was used to extract each peptide coding sequence, and 1:1 orthologs (n=7,154) between *Z. japonica* and *Z. marina* were identified using best reciprocal BLAT hits. The 4DTv rate among orthologs was calculated and estimated using 10,000 bootstrap estimates (0.0795-0.0816; 90% CI). Applying the WGD age to the 4DTv neutral rate, the divergence time between *Z. japonica* and *Z. marina* was estimated to be between 9.86 - 12.67 MYA, which is consistent with the upper bound estimate of 11.9 mya from Coyer et al. (2013).

#### Estimating the mutation rate

The divergence time between *Z. japonica* and *Z. marina* was represented as a lognormal distribution (Mean: 11.1542 MYA; SD: 0.07). An estimate of a neutral mutation rate was calculated based on the synonymous mutation rate ($K_s$) among *Z. japonica* orthologs and four *Z. marina* populations (JS, BB, NC, and FR). The median Ks peak between *Z. japonica* and *Z. marina* was approximately 0.211, which was consistent across all 4 populations. The neutral mutation rate was calculated as $r = K_s / (2 * \text{divergence age}) = 9.462 * 10^{-9}$ per generation per year.

#### SNP calling and filtering including *Z. japonica*

One *Z. japonica* sample was included for genetic variant calling. The GVCF format file for *Z. japonica* was generated in the same way with the *Z. marina*. All the GVCF files (190 *Z. marina* samples + 1 *Z. japonica*) were combined by CombineGVCFs (GATK4). GenotypeGVCFs (GATK4) was used to call genetic variants. BCFtools was used to remove SNPs within 20 base pairs of an indel or other variant type. Then we extracted only SNPs (10,562,762 SNPs), and excluded the other types of variants. VariantsToTable (GATK4) was used to extract INFO annotations. SNPs meeting one or more than one of the following criteria were marked by VariantFiltration (GATK4): MQ < 40.0; FS > 60.0; QD < 10.0; MQRandSum > 2.5 or MQRandSum < -2.5; ReadPosRandSum < -2.5; ReadPosRandSum > 2.5; SOR > 3.0; DP > 11185.0 (2 * average DP). Those SNPs were excluded by SelectVariants (GATK4). A total of 4,873,274 SNPs were retained. VCFtools was used to convert individual genotypes to missing data when GQ < 30 or DP < 10. Individual homozygous reference calls with one or more read supporting the variant allele, and individual homozygous variant calls supporting the reference allele, were also converted to missing data using a custom Python3 script. Only bi-allelic SNPs were kept (4,775,984 SNPs). To avoid the reference-related biases, we focused on the core genes shared by all individual samples. Bedtools was used to find overlap between the SNPs and the core genes in the six main chromosomes (18,717 genes), and only those SNPs were kept (1,483,603 SNPs). Thirty-seven ramets were excluded from ZM_HQ_SNPs, including 10 ramets related to geitonogamous (within-clone) selfing (Supplementary Table 2), 17 ramets representing replicated genotypes (Supplementary Table 3), and 10 ramets with high missing rate (Supplementary Fig. 1). After excluding these ramets, some SNPs became monomorphic, which were then excluded. SnpEff was used to annotate each SNP. To obtain putatively neutral SNPs, we only kept SNPs annotated as "synonymous_variant" (ZMZJ_Neutral_SNPs, 171,756 SNPs). Then only SNPs without any missing data were kept (ZMZJ_D_SNPs, 20,341 SNPs).

Coyer, J. A., Hoarau, G., Kuo, J., Tronholm, A., Veldsink, J., & Olsen, J. L. (2013). Phylogeny and temporal divergence of the seagrass family Zosteraceae using one nuclear and three chloroplast loci. Systematics and Biodiversity, 11(3), 271-284.

Keilwagen, J., Hartung, F., & Grau, J. (2019). GeMoMa: Homology-Based Gene Prediction Utilizing Intron Position Conservation and RNA-seq Data. Methods in molecular biology (Clifton, NJ), 1962, 161-177.

Olsen, J. L., Rouzé, P., Verhelst, B., Lin, Y. C., Bayer, T., Collen, J., ... & Van de Peer, Y. (2016). The genome of the seagrass *Zostera marina* reveals angiosperm adaptation to the sea. Nature, 530(7590), 331-335.

Pertea, G., & Pertea, M. (2020). GFF utilities: GffRead and GffCompare. F1000Research, 9.

Schmutz, J., & Lovell, J. (2021). GENESPACE R Package (GENESPACE) v1. 0 (No. GENESPACE R Package). HudsonAlpha Institute for Biotechnology; Lawrence Berkeley National Lab. (LBNL), Berkeley, CA (United States).

### Supplementary Note 3 - divergence time estimation using ASTRAL

#### Constructing an ASTRAL species tree

Illumina sequencing libraries from 190 samples of *Zostera marina* were cleaned (Illumina reads less than 75 bp after trimming for adapter and quality (q < 20) were removed) and de novo assembled using HipMer (k=51). To build suitable gene trees (and a subsequent species tree), transcript sequences from *Z. marina* (v3.1, n = 21,483) were aligned against each HipMer (Georganas et al. 2015) assembly (n=212) using BLAT (v30) (Kent 2002). To find conserved gene sequences in all samples, a transcript was considered 'present' if it had (i) a single alignment greater than 60% identity and coverage; or (ii) a transcript was covered at least to 85% by 3 or fewer alignments among contigs with identity >85%. Next, two quality control assessments were performed prior to gene tree analysis. First, using samples identified as clones, 1016 genes were found to be inconsistently aligned (considered present in one clone but absent in another) and were removed from consideration. Clones were then removed from the dataset, retaining only one representative sample per clone set. Secondly, the total number of genes considered present was counted within all samples, excluding libraries with counts fewer than 17,500 as those were considered fragmented and of too low quality. The final dataset included 129 samples and 20,100 aligned transcripts, comprising 18,311 genes present in 97.9% of all accessions (core gene set). CDS and protein sequences from each transcript alignment were predicted using GeMoMa (v1.7; Keilwagen et al. 2019) using constrained search where each transcript's best alignment location was extracted using bedtools (Quinlan 2014) (getfasta) with 800 bp buffer. A random subset of 617 genes was then selected for building gene trees and the final ASTRAL tree. Admixed samples (populations Washington State and Bodega Bay; discussed in main ms) were not included for gene tree construction. Gene trees were constructed by first aligning CDS sequences together using MAFFT (v7.475; Katoh et al. 2002) (parameters: mafft --localpair --phylipout --maxiterate 1000), then generating individual gene trees with IQTREE (v2.1.2) (Nguyen et al. 2015; parameters -B 1000 -m K2P -T auto). All gene trees were used jointly as input for a species tree analysis with ASTRAL (Zhang et al. 2018) v5.7.3, using a map file to join multiple samples representing the same population.

#### Divergence time estimates using StarBEAST2

To investigate the divergence among *Z. marina* populations, the gene alignments (generated by MAFFT as outlined in the ASTRAL species tree section) were analyzed using StarBEAST2 (v2.6.3) (Bouckaert et al. 2014; Heled and Drummond 2010). As age calibration, the divergence between *Z. japonica* and *Z. marina* was estimated using the WGD event that occurred approximately 67 Mya (Olsen et al. 2016) (Supplementary Note 2). Next, protein sequences were predicted from an assembly of *Zostera japonica* (Xiaomei Zhang pers. comm., available at doi.org/10.6084/m9.figshare.21626327.v1, and Zhang et al. 2019) using the GeMoMa pipeline outlined above. GeMoMa predicted 8,687 peptide sequences that, when aligned to *Z. marina* peptides using BLAT [v30] (Kent 2002), yielding 7154 reciprocal best orthologs. 4DTV rate calculation among best hit orthologs found a clear peak estimated between 0.0795- 0.0816 (90% CI-from 10,000 bootstrap estimates). Based on the neutral divergence among syntenic paralogs from the *Zostera* WGD event, we obtained a divergence estimate between *Z. japonica* and *Z. marina* at 9.86 - 12.67 mya, consistent with the upper bound of divergence times provided from Coyer et al. (11.8-2.9 mya; Coyer et al. 2013). A median age of 11.01 mya was selected for subsequent StarBEAST2 anaylsis.

The xml file was generated from MAFFT alignments (phylip format) using seqmagick (v0.6.2; Shen et al. 2016) to convert phylip format to nexus. The XML file required for StarBEAST2 was generated using BEAUTi from 70 randomly selected alignments with no missing data in a subset of populations (four samples represented per population): ALS, JN, JS, SD, MA, NN, SW, FR, with *Z. japonica* and *Z. marina* reference sequences. Parameters for the StarBEAST2 run were as follows: gene ploidy = 2; constant population sizes; population size parameter = 0.03 (which assumes a 10K effective population size and a generation time of 3 years); Gamma site model with estimated substitution rate; HKY substitution model; estimate kappa; empirical nucleotide frequencies; strict molecular clock; estimated clock rate; Yule model; Outgroup= *Z. japonica* ; Outgroup divergence constrained with a lognormal prior [M=11.01; S=0.01; mean in real space, use originate]; MCMC chain length = 200,000,000; store every 200,000; pre-burnin = 0. starBEAST was run in triplicate, inspecting each run with Tracer to check for model convergence (Barido-Sottani et al. 2018) (20% burn-in; Effective sample sizes [ESS]>300). TreeAnnotator was to summarize each run (20% burn-in, median peak height, 0.5 posterior probability limit) which were then combined using LogCombiner. Finally, the log combined tree file was re-summarized with TreeAnnotator using the parameters listed above. Note that An ASTRAL/StarBEAST2 analysis requires an estimate of generation time and population size, both of which are subject to large error in eelgrass owing to large differences in clonal vs. sexual reproduction among populations (see also Supplementary Table 3).

Barido-Sottani, J., Bošková, V., Plessis, L. D., Kühnert, D., Magnus, C., Mitov, V., Müller, N. F., PecErska, J., Rasmussen, D. A., Zhang, C., Drummond, A. J., Heath, T. A., Pybus, O. G., Vaughan, T. G., & Stadler, T. (2018). Taming the BEAST-A Community Teaching Material Resource for BEAST 2. Systematic Biology, 67(1), 170–174.

Bouckaert, R., Heled, J., Kühnert, D., Vaughan, T., Wu, C.-H., Xie, D., Suchard, M. A., Rambaut, A., & Drummond, A. J. (2014). BEAST 2: a software platform for Bayesian evolutionary analysis. PLoS Computational Biology, 10(4), e1003537.

Coyer, J. A., Hoarau, G., Kuo, J., Tronholm, A., Veldsink, J., & Olsen, J. L. (2013). Phylogeny and temporal divergence of the seagrass family Zosteraceae using one nuclear and three chloroplast loci. Systematics and Biodiversity, 11(3), 271–284.

Georganas, E., Buluç, A., Chapman, J., Hofmeyr, S., Aluru, C., Egan, R., Oliker, L., Rokhsar, D., & Yelick, K. (2015). HipMer: an extreme-scale de novo genome assembler. Proceedings of the International Conference for High Performance Computing, Networking, 1–11.

Heled, J., & Drummond, A. J. (2010). Bayesian inference of species trees from multilocus data. Molecular Biology and Evolution, 27(3), 570–580.

Katoh, K., Misawa, K., Kuma, K.-I., & Miyata, T. (2002). MAFFT: a novel method for rapid multiple sequence alignment based on fast Fourier transform. Nucleic Acids Research, 30(14), 3059–3066.

Keilwagen, J., Hartung, F., & Grau, J. (2019). GeMoMa: Homology-Based Gene Prediction Utilizing Intron Position Conservation and RNA-seq Data. Methods in Molecular Biology, 1962, 161–177.

Kent, W. J. (2002). BLAT — The BLAST -Like Alignment Tool. Genome Research, 12, 656–664.

Lovell, J. T., Sreedasyam, A., Schranz, M. E., Wilson, M., Carlson, J. W., Harkess, A., Emms, D., Goodstein, D. M., & Schmutz, J. (2022). GENESPACE tracks regions of interest and gene copy number variation across multiple genomes. eLife, 11. https://doi.org/10.7554/eLife.78526

Nguyen, L.-T., Schmidt, H. A., von Haeseler, A., & Minh, B. Q. (2015). IQ-TREE: a fast and effective stochastic algorithm for estimating maximum-likelihood phylogenies. Molecular Biology and Evolution, 32(1), 268–274.

Olsen, J. L., Rouzé, P., Verhelst, B., Lin, Y. C., Bayer, T., Collen, J., Dattolo, E., De Paoli, E., Dittami, S., Maumus, F., Michel, G., Kersting, A., Lauritano, C., Lohaus, R., Töpel, M., Tonon, T., Vanneste, K., Amirebrahimi, M., Brakel, J., … Van De Peer, Y. (2016). The genome of the seagrass *Zostera marina* reveals angiosperm adaptation to the sea. Nature, 530(7590), 331–335.

Quinlan, A. R. (2014). BEDTools: The Swiss-army tool for genome feature analysis. Et Al [Current Protocols in Bioinformatics], 47(1), 11.12.1–34.
Shen, W., Le, S., Li, Y., & Hu, F. (2016). SeqKit: A Cross-Platform and Ultrafast Toolkit for FASTA/Q File Manipulation. PloS One, 11(10), e0163962.
Zhang, C., Rabiee, M., Sayyari, E., & Mirarab, S. (2018). ASTRAL-III: Polynomial time species tree reconstruction from partially resolved gene trees. BMC Bioinformatics, 19(Suppl 6), 15–30.
Zhang X, Zhou Y, Li Y-L, Liu J-X (2019) Development of microsatellite markers for the seagrass *Zostera japonica* using next-generation sequencing. Molecular Biology Reports 46: 1335-1341 doi 10.1007/s11033-018-4491-2

### Supplementary Note 4 - using the cpDNA haplotype clock as an alternative dating estimate

#### Estimating chloroplast haplotype divergence time

We used chloroplast synonymous substitutions to validate the divergence time between the SD and BB populations and the rest of the Pacific. The shortest synonymous mutation distance ($D_{Ss}$) between the two groups is 10 mutations, for example, between SD10 and WAS03. The longest distance ($D_{Sl}$) is 13 mutation steps, for example, between haplotypes represented in SD11 and JS03. The total of 9731 synonymous sites in chloroplast protein-coding genes (S) was calculated based on a custom-made annotation. The chloroplast synonymous substitution rate ($K_S$) range of 1.1 – 2.9 x $10^{-9}$ substitutions per site per year was obtained from Muse (2000). The divergence time of the Californian and the main Pacific chloroplasts is expected to fall between the fastest and the slowest evolution scenario, which are $D_{Ss} / (2 * K_{Smax} * S) = 0.177$ Mya and $D_{Sl} / (2 * K_{Smin} * S) = 0.607$ Mya correspondingly. Thereby the average expected divergence time based on chloroplast synonymous substitutions is 0.392 Mya which is in accordance to the nuclear-based calculated divergence time (0.3525 Mya).

Muse SV. 2000. Examining rates and patterns of nucleotide substitution in plants. In: Doyle JJ, Gaut BS, editors. Plant Molecular Evolution. Dordrecht: Springer Netherlands. p. 25–43. Available from: http://link.springer.com/10.1007/978-94-011-4221-2_2

### Supplementary Note 5 - *Zostera marina* and *Z. pacifica* in the San Diego to Baja region

Our data raise the question whether the deep divergent San Diego SD lineage of *Z. marina* may in fact be the newly described *Z. pacifica*. The presence of a cryptic species of *Zostera* in the California Channel Islands was confirmed in an extensive genetic survey of the California Bight based on nine microsatellite loci (Coyer et al. 2008 and review therein). *Zostera pacifica* (wide-leaved eelgrass) was found to be restricted to the California Channel Islands and the adjacent mainland and parts of South San Diego Bay. The problem is that Z. *pacifica* is typically wide-leafed but *Z. marina* can also be wide-leafed, rendering this morphological trait unreliable. Rather, the only conclusive diagnostic feature is molecular and entails the failure of microsatellite locus CT-20 (Reusch 2000) to amplify in *Z. pacifica* (Coyer et al. 2008). A STRUCTURE analysis further revealed varying degrees of introgression between the two species at three locations. Coyer et al. (2008) concluded that the distribution of *Z. pacifica* follows the glacial age land mass of the California Bight. The recognition of two species is important in terms of protection of biodiversity, transplant mitigation and management, which has been extensive in southern California (Coyer et al. 2008, Olsen et al. 2014).

The 12 San Diego (SD) samples used in the present study amplified successfully with the CT-20 microsatellite locus, thus designating them as *Z. marina*. However, our resequencing data from these samples reveals the split between SD and BB (Fig. 2G), as well as introgression of WAS and BB with SD (Fig. 2D) in a phylogeographically shallow timeframe. This begs the age-old question of how much divergence is needed to recognize a species vs. a subspecies, variety or race? At present, we continue to designate these samples as intra-specific admixtures of *Z. marina,* but noting that recognition of an inter-specific admixture would not affect any of our conclusions in this paper. What will be interesting are the potential differences in functional gene traits that population genomics makes possible. With this in mind, a second, high-quality reference genome of a non-admixed E Pacific *Z. marina* is currently being sequenced as part of our characterization of the *Z. marina* pangenome; a companion to the currently available, chromosomal-level reference genome originating in the North-East Atlantic (Baltic Sea; Olsen et al. 2016; Ma et al. 2021).

Coyer, J.A., Miller, K.A., Engle, J.M., Veldsink, J., Cabello-Pasini, A., Stam W.T. & Olsen, J.L 2008. Eelgrass meadows in the California Channel Islands and adjacent coast reveal a mosaic of two species, evidence for introgression and variable clonality. Annals Bot. 101:73-87.

Ma X, Olsen JL, Reusch TBH, Procaccini G, Kudrna D, Williams M, Grimwood J, Rajasekar S, Jenkins J, Schmutz J, Van de Peer Y (2021) Improved chromosome-level genome assembly and annotation of the seagrass, *Zostera marina* (eelgrass) F1000Research 10: 289

Olsen, J.L., Coyer, J.A., Chesney, B. 2014. Numerous mitigation transplants of eelgrass *Zostera marina* in southern California shuffle genetic diversity and may promote hybridization with *Z. pacifica.* Biological Conservation 176: 133-143

Reusch TBH (2000) Five microsatellite loci in eelgrass *Zostera marina* and a test of cross-species amplification in *Z. noltii* and *Z. japonica*. Molecular Ecology 9: 371-373

### Supplementary Tables

#### Supplementary Table 1: Population metadata for 16 worldwide *Zostera marina* populations

| | Region | Population code | Population location | Latitude | Longitude | *ZEN Cross ref | Sample Numbering |
|---|---|---|---|---|---|---|---|
| West Pacific | | | | | | | |
| | Japan North | JN | Akkeshi-ko Estuary, Hokkaido, Japan | 43.021N | 144.903E | JN-A | JN101–JN112 |
| | Japan South | JS | Ikunoshima, Japan (Inland sea) | 34.298N | 132.916E | JS-A | JS201–JS 212 |
| East Pacific | | | | | | | |
| | Bering Sea **LME | ASL | Safety Lagoon, Alaska, USA | 64.485N | 164.762W | Non-ZEN | ASL1–ASL12 |
| | Gulf of Alaska LME | ALI | Izembek Lagoon, Alaska, USA | 55.329N | 162.821W | Non-ZEN | ALI301–ALI312 |
| | California Current LME | WAS | Willapa Bay, Washington State, USA | 46.474N | 124.028W | WAS-A | WAS401–WAS412 |
| | California Current LME | BB | Westside Park, Bodega Bay, California, USA | 38.320N | 123.055W | BB-A | BB501–BB512 |
| | California Current LME | SD | Shelter Island, San Diego Bay, California, USA | 32.714N | 117.225W | SD-A | SD601–SD612 |
| West Atlantic | | | | | | | |
| | Quebec | QU | Pointe-Lebel, Quebec, Canada | 49.112N | 68.176W | QU-A | QU701–QU711 |
| | Massachusetts | MA | Dorothy Cove, Massachusetts, USA | 42.420N | 70.915W | MA-A | MA801–MA812 |
| | North Carolina | NC | Middle Marsh, North Carolina, USA | 34.692N | 76.623W | NC-A | NC 1103–NC 1112 |
| East Atlantic | | | | | | | |
| | Northern Norway | NN | Røvika, Norway (near Bødo) | 67.268N | 15.257E | NN-B | NN1201–NN1212 |
| | Sweden | SW | Torserød, Sweden (west coast) | 58.313N | 11.549E | SW-A | SW1401–SW1412 |
| | Wales | WN | Port Dinllaen, Wales, UK | 52.991N | 4.450W | WN-A | WN1501–WN1512 |
| | Portugal | PO | Culatatra, Ria Formosa, S. Portugal | 37.040N | 7.910W | PO-A | PO1601–PO1611 |
| Mediterranean Sea | | | | | | | |
| | France | FR | Bouzigues, Thau Lagoon, France | 43.447N | 3.662E | FR-A | FR1701–FR1712 |
| | Croatia | CZ | Adriatic Sea, Posedarje, Croatia | 44.212N | 15.491E | CR-A | CZ1801–CZ1812 |

*The *Zostera* Experimental Network (ZEN) has produced extensive ecological and physical metadata for 50 populations of which the above represent 16 of them. All data used in their analyses, and associated R code, are available at https://doi.org/10.5281/zenodo.6808753 with the exception of the genetic data, available at https://doi.org/10.5281/zenodo.3660013 . **LME = large marine ecosystem

#### Supplementary Table 2: Possible parent-descendant pairs under self-fertilization as detected using the shared heterozygosity index.

| Selfing Pair | Parent Ramet | Descendant Ramet |
|---|---|---|
| SP_01 | NN05 | NN02 |
| SP_02 | NN05 | NN06 |
| SP_03 | NN05 | NN07 |
| SP_04 | NN05 | NN09 |
| SP_05 | NN05 | NN10 |
| SP_06 | NN08 | NN02 |
| SP_07 | NN08 | NN06 |
| SP_08 | NN08 | NN07 |
| SP_09 | NN08 | NN09 |
| SP_10 | NN08 | NN10 |
| SP_11 | SD03 | SD02 |
| SP_12 | SD03 | SD12 |
| SP_13 | SD12 | SD02 |
| SP_14 | WN02 | WN03 |
| SP_15 | WN02 | WN07 |
| SP_16 | WN12 | WN05 |

See Supplementary Fig. 3 for further details on the shared heterozygosity index.

#### Supplementary Table 3: Clone mates detected based on the shared heterozygosity index.

| Genet | Clonemates | | | | | | |
|---|---|---|---|---|---|---|---|
| Genet_01 | BB04 | BB05 | | | | | |
| Genet_02 | BB09 | BB10 | | | | | |
| Genet_03 | JS03 | JS04 | | | | | |
| Genet_04 | NN02 | NN06 | NN07 | NN09 | NN10 | | |
| Genet_05 | PO02 | PO05 | PO07 | PO08 | PO10 | PO11 | PO12 |
| Genet_06 | PO03 | PO04 | PO06 | PO09 | | | |
| Genet_07 | SD04 | SD11 | | | | | |
| Genet_08 | SD06 | SD09 | | | | | |
| Genet_09 | WN04 | WN09 | | | | | |
| Genet_10 | WN06 | WN10 | | | | | |
| Genet_11 | NN05 | NN08 | | | | | |

See Supplementary Fig. 3 for further details on the shared heterozygosity index.

#### Supplementary Table 4: Statistical comparison of genetic diversity measures

Genetic diversity measures ($\pi$, $H_{OBS}$) were subject to 1-way analysis of variance, followed by Tukey Cramer HSD post-hoc test, using a family-wise error rate of $\alpha$=0.05. Different letters indicate statistically different mean values. Population abbreviations are given in Fig. 1.

| Population | | phi | Population | | Hobs |
|---|---|---|---|---|---|
| JS | A | 0.0816 | JS | A | 0.0984 |
| JN | B | 0.0695 | JN | B | 0.0852 |
| BB | C | 0.0584 | BB | B | 0.0798 |
| WAS | D | 0.0450 | WAS | C | 0.0596 |
| SD | E | 0.0302 | SD | D | 0.0421 |
| NC | F | 0.0148 | ASL | E | 0.0210 |
| ALI | F G | 0.0133 | ALI | E F | 0.0203 |
| ASL | F G H | 0.0116 | NC | E F | 0.0152 |
| PO | F G H | 0.0113 | PO | E F G | 0.0123 |
| CZ | F G H I | 0.0096 | FR | F G | 0.0116 |
| FR | F G H I | 0.0094 | CZ | F G | 0.0107 |
| MA | F G H I | 0.0078 | MA | F G | 0.0090 |
| SW | G H I | 0.0064 | SW | F G | 0.0090 |
| WN | H I | 0.0053 | WN | F G | 0.0088 |
| QU | H I | 0.0047 | QU | G | 0.0057 |
| NN | I | 0.0025 | NN | G | 0.0028 |

#### Supplementary Table 5: Matrix of $F_{ST}$-values among population pairs

| **Pop\Pop** | **JN** | **JS** | **ASL** | **ALI** | **WAS** | **BB** | **SD** | QU | MA | NC | NN | SW | WN | PO | FR | CZ |
|---|---|---|---|---|---|---|---|---|---|---|---|---|---|---|---|---|
| **JN** | 0.0 | 0.523 | 0.399 | 0.31 | 0.474 | 0.525 | 0.692 | 0.629 | 0.635 | 0.616 | 0.58 | 0.634 | 0.587 | 0.518 | 0.64 | 0.633 |
| **JS** | | 0.0 | 0.698 | 0.618 | 0.649 | 0.641 | 0.724 | 0.731 | 0.738 | 0.723 | 0.69 | 0.74 | 0.697 | 0.628 | 0.739 | 0.736 |
| **ASL** | | | 0.0 | 0.503 | 0.613 | 0.686 | 0.881 | 0.898 | 0.885 | 0.851 | 0.897 | 0.89 | 0.889 | 0.878 | 0.881 | 0.877 |
| **ALI** | | | | 0.0 | 0.498 | 0.59 | 0.852 | 0.916 | 0.896 | 0.845 | 0.923 | 0.904 | 0.908 | 0.876 | 0.887 | 0.882 |
| **WAS** | | | | | 0.0 | 0.389 | 0.704 | 0.782 | 0.783 | 0.761 | 0.753 | 0.785 | 0.755 | 0.708 | 0.783 | 0.779 |
| **BB** | | | | | | 0.0 | 0.556 | 0.782 | 0.784 | 0.764 | 0.748 | 0.787 | 0.752 | 0.69 | 0.784 | 0.78 |
| **SD** | | | | | | | 0.0 | 0.909 | 0.904 | 0.881 | 0.897 | 0.908 | 0.895 | 0.857 | 0.9 | 0.898 |
| QU | | | | | | | | 0.0 | 0.391 | 0.358 | 0.594 | 0.445 | 0.473 | 0.738 | 0.609 | 0.57 |
| MA | | | | | | | | | 0.0 | 0.229 | 0.457 | 0.328 | 0.345 | 0.644 | 0.539 | 0.498 |
| NC | | | | | | | | | | 0.0 | 0.345 | 0.277 | 0.28 | 0.502 | 0.464 | 0.427 |
| NN | | | | | | | | | | | 0.0 | 0.26 | 0.401 | 0.792 | 0.587 | 0.545 |
| SW | | | | | | | | | | | | 0.0 | 0.175 | 0.658 | 0.517 | 0.455 |
| WN | | | | | | | | | | | | | 0.0 | 0.704 | 0.54 | 0.495 |
| PO | | | | | | | | | | | | | | 0.0 | 0.601 | 0.609 |
| FR | | | | | | | | | | | | | | | 0.0 | 0.503 |
| CZ | | | | | | | | | | | | | | | | 0.0 |

Population abbreviations see Fig. 1. Values are calculated based on the core SNP set (11,705 SNP, Supplementary Fig. 1). Pacific populations are shown in boldface. All pairwise $F_{ST}$ have significant $p$ values ($p < 0.001$), based on 1,000 permutation runs, hence only the half diagonal matrix is depicted. Note that JN shows the smallest $F_{ST}$ of all Pacific populations with any other population from the Atlantic side (Atlantic + Mediterranean Sea).

### Supplementary Figures

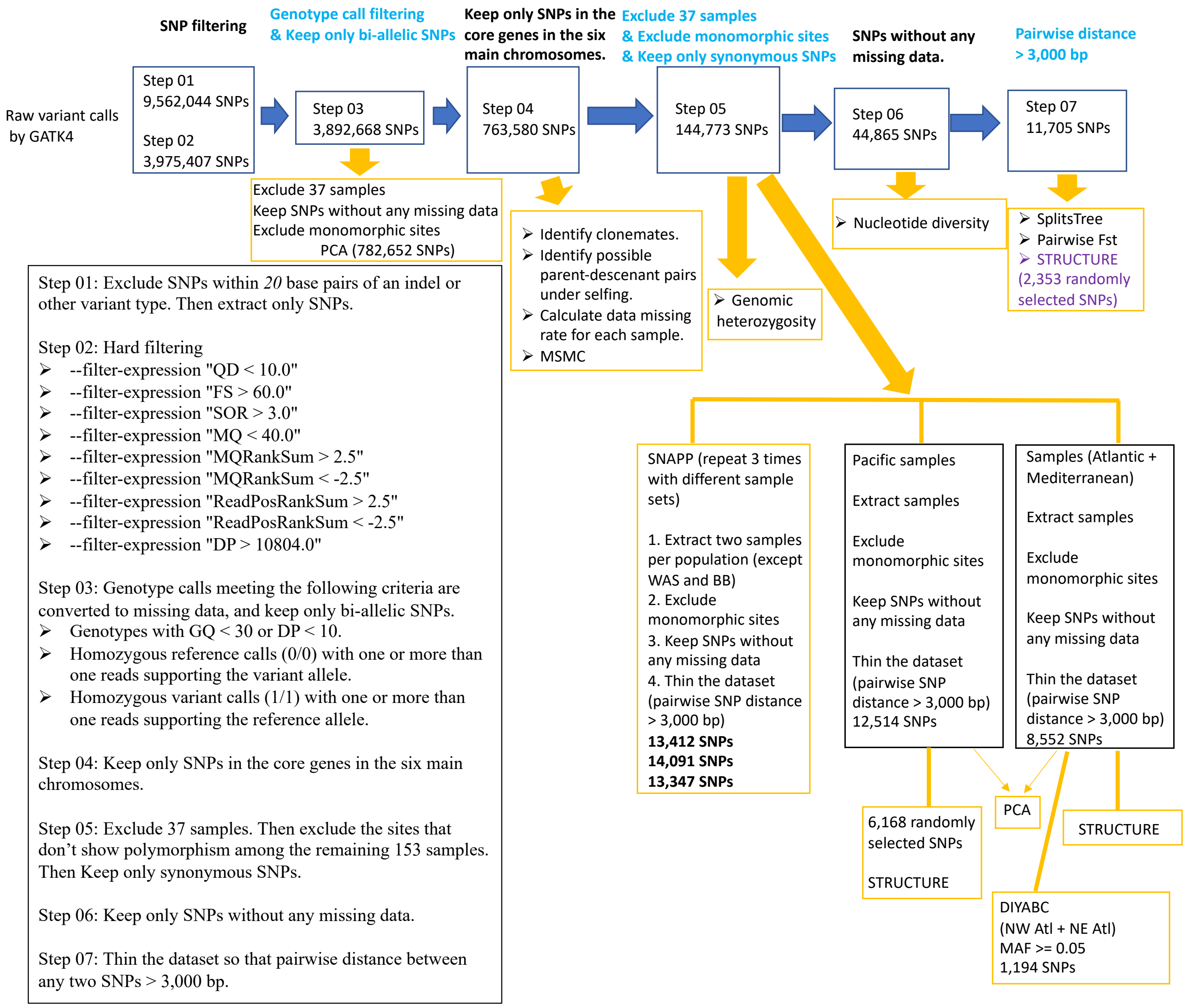

#### Supplementary Fig. 1 | Workflow for SNP calling and filtering

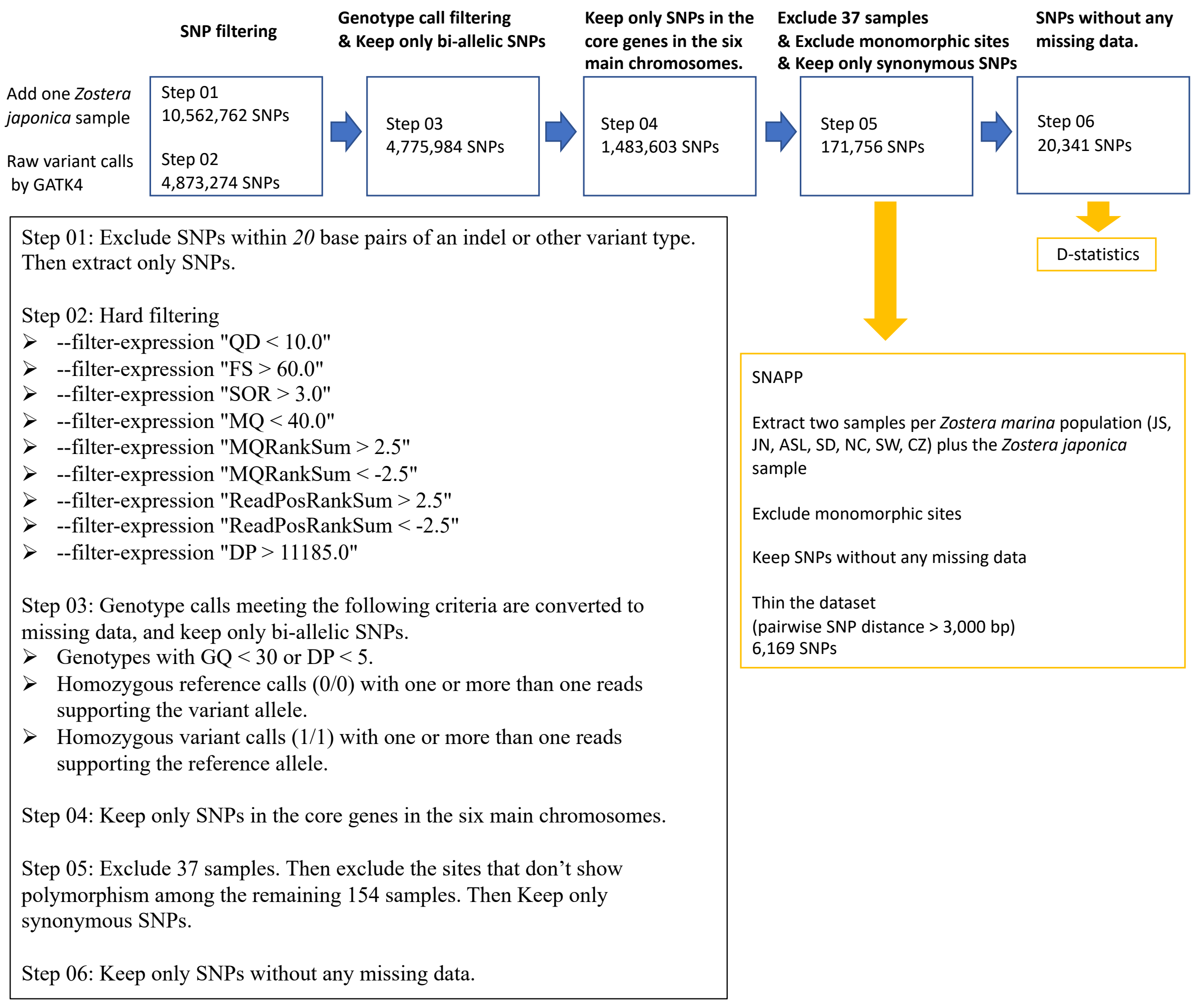

**Supplementary Fig. 2 | Workflow for SNP calling and filtering that required an outgroup.**

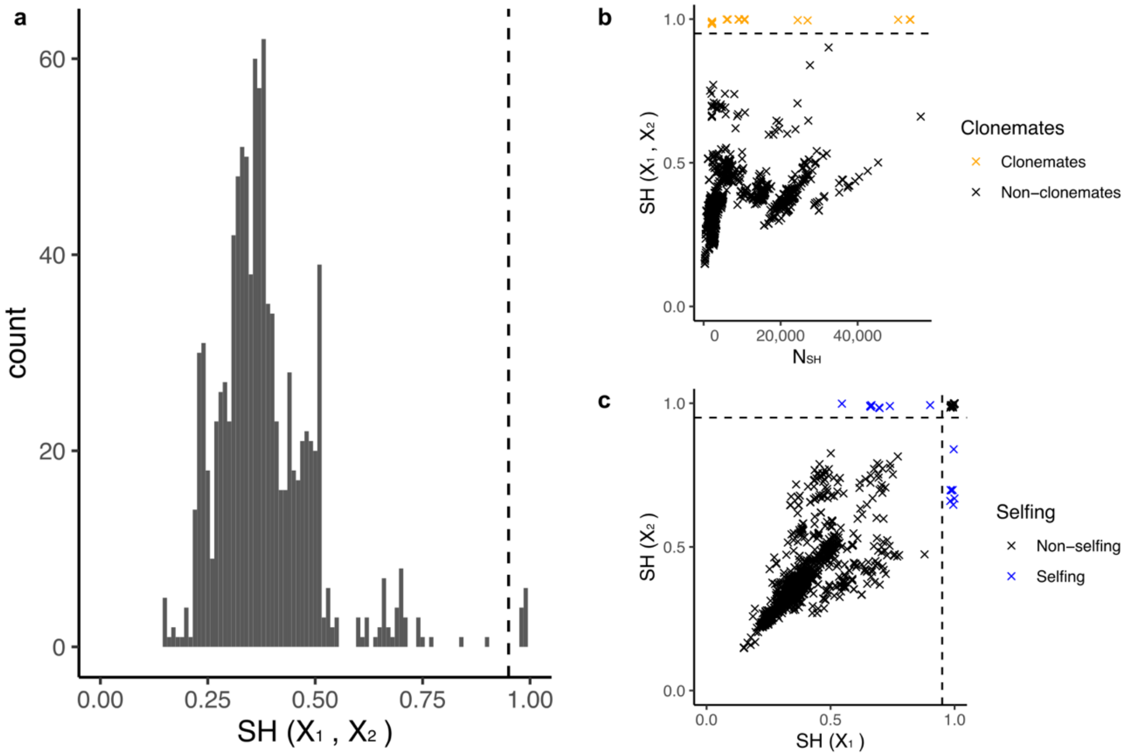

**Supplementary Fig. 3 | Detecting clone mates and possible parent-descendant pairs under selfing based on shared heterozygosity (SH)** according to Yu et al. (2022) a**,** Histogram of pairwise SH($X_1$, $X_2$) for each pair of ramets within the same population. The threshold is set to 0.95, indicated by the vertical dashed line. **b,** Identification of pairs of ramets belonging to the same genet (=clone mates) based on SH($X_1$, $X_2$). Clone mate pairs show SH($X_1$, $X_2$) > 0.95. **c,** Identification of possible parent-descendant pairs under selfing based on SH($X_1$) and SH($X^2$). Ramets originating via selfing inherit a subset of the heterozygous loci from the parent.

Yu L, Stachowicz JJ, DuBois K, Reusch TBH (2022) Detecting clonemate pairs in multicellular diploid clonal species based on a shared heterozygosity index. Mol Ecol Resources online early: https://doi.org/10.1111/1755-0998.13736 doi 10.1101/2022.02.16.480681

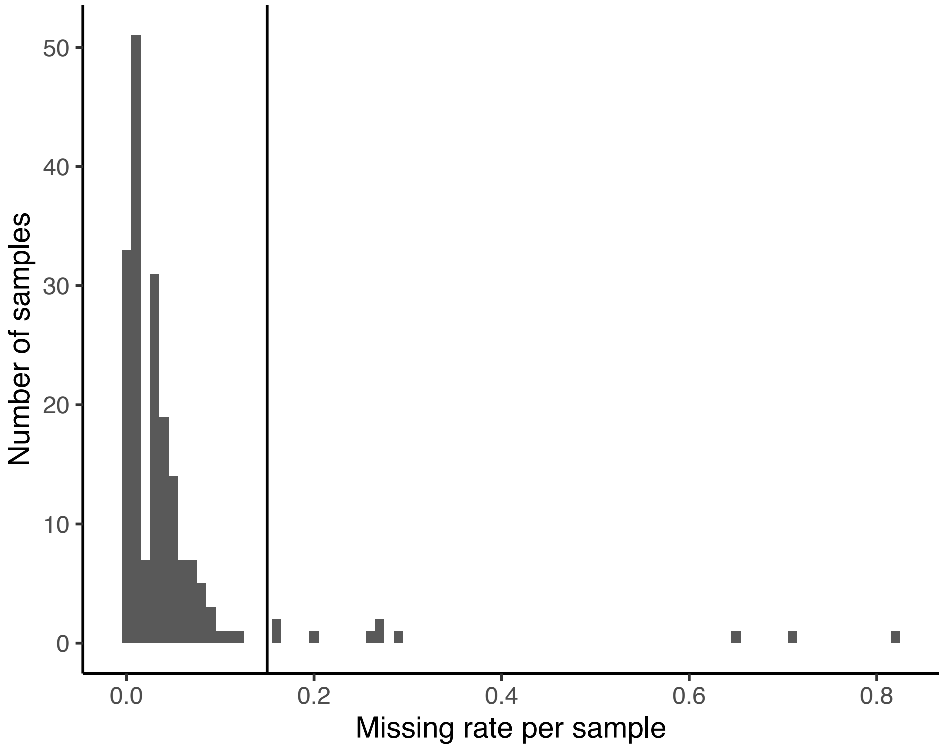

**Supplementary Fig. 4 | Missing data distribution for each sample.** The data missing rate for each sample was calculated based on SNP data set ZM_HQ_SNPs (763,580 SNPs). Missing rate=number of loci with missing data/763,580. Samples with <15% missing data were retained. Ten of the 190 samples had a missing rate >15% (ALI01, ALI02, ALI03, ALI04, ALI05, ALI06, ALI10, ALI16, QU03 and SD08) and were excluded from the data set.

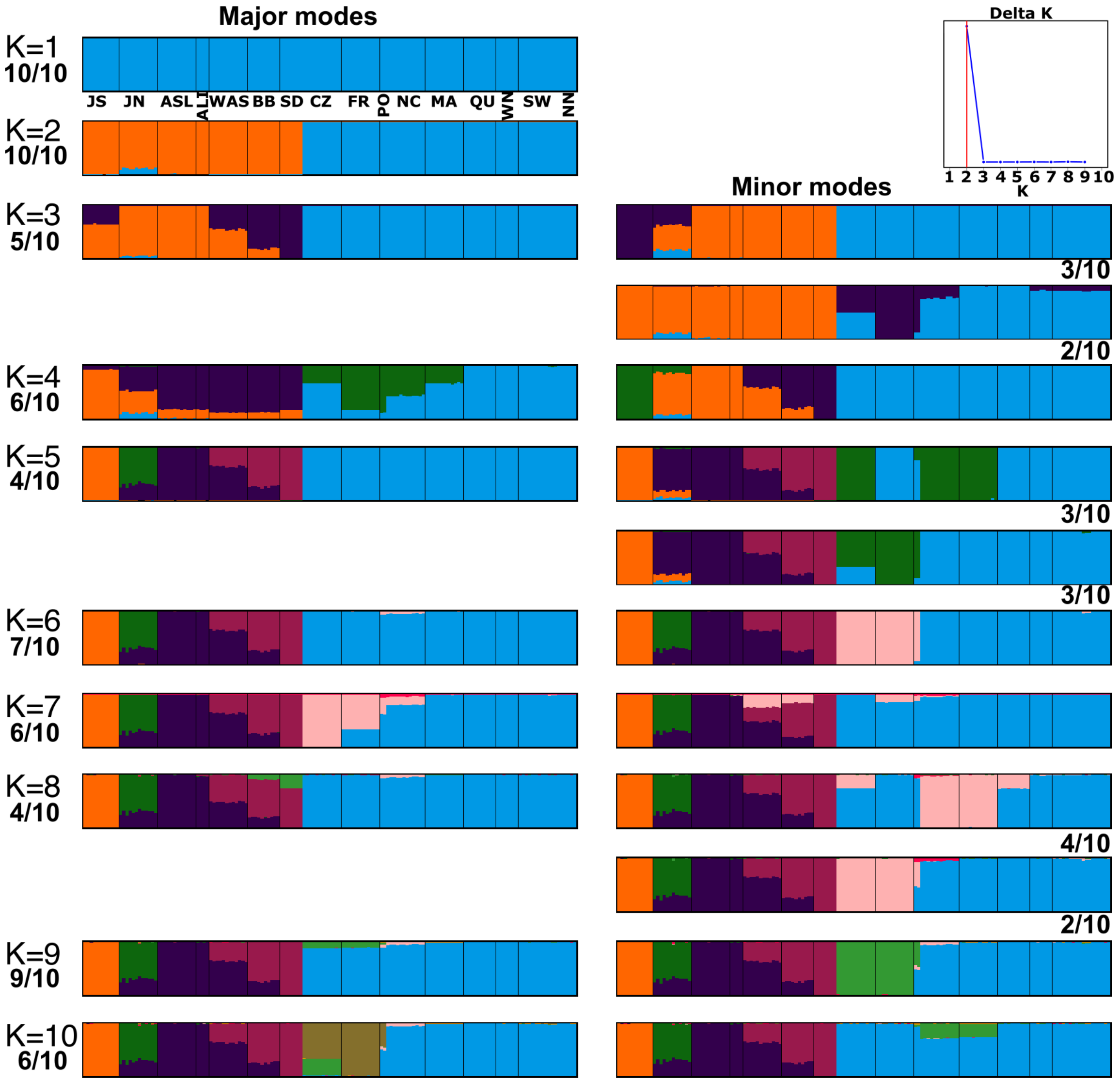

**Supplementary Fig. 5 | Global STRUCTURE results for K from 1 to 10.** The analysis was repeated 10 times with the same parameters. The number indicates how many times the mode occurred among the 10 runs.

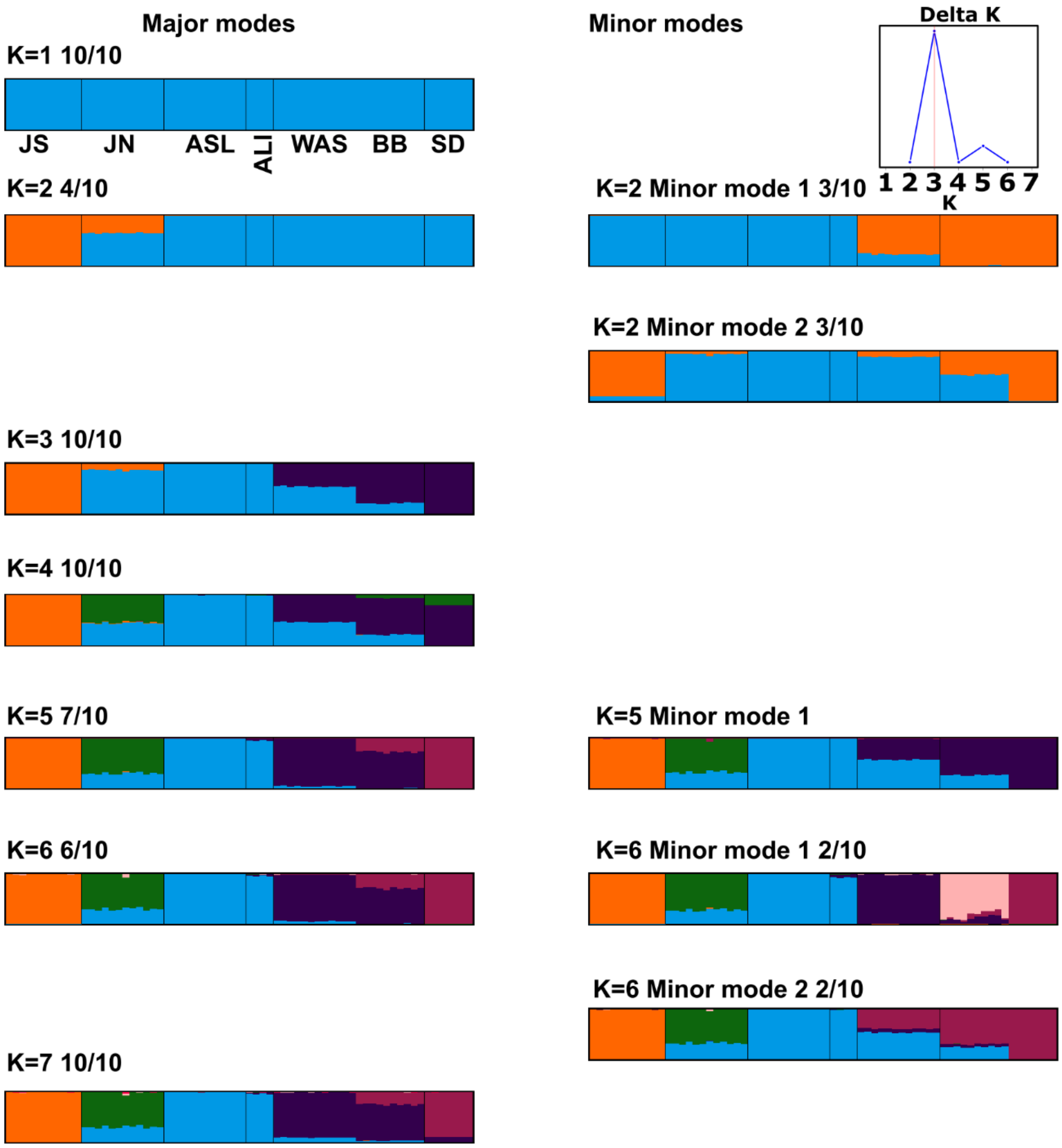

**Supplementary Fig. 6 | STRUCTURE results for Pacific populations only for K from 1 to 7.** The analysis was repeated 10 times with the same parameters. The number indicates how many times the mode occurred among runs.

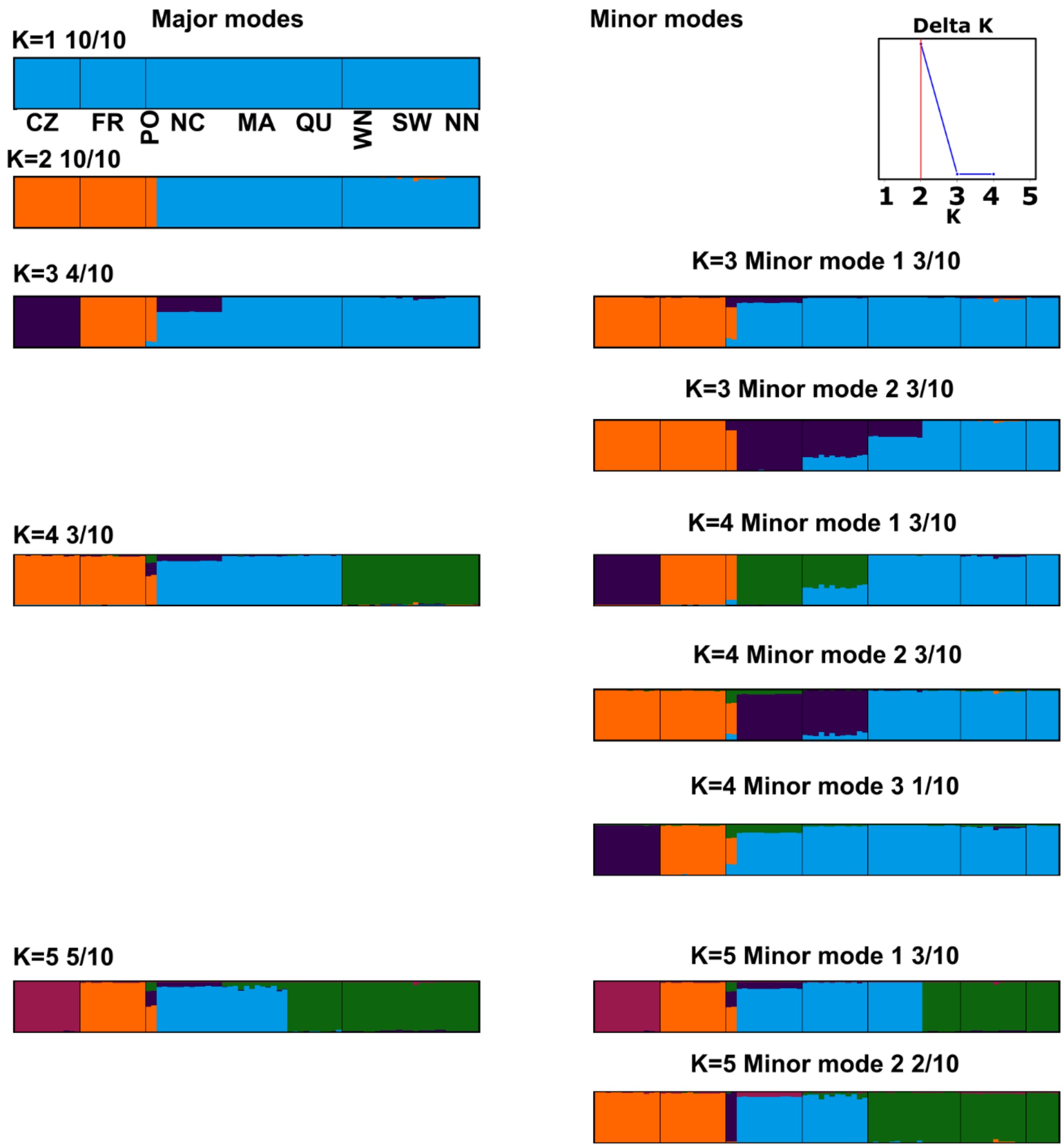

**Supplementary Fig. 7 | STRUCTURE results for Atlantic populations for K from 1 to 5.**

The analysis was repeated 10 times with the same parameters. The number indicates how many times the mode occurred among runs.

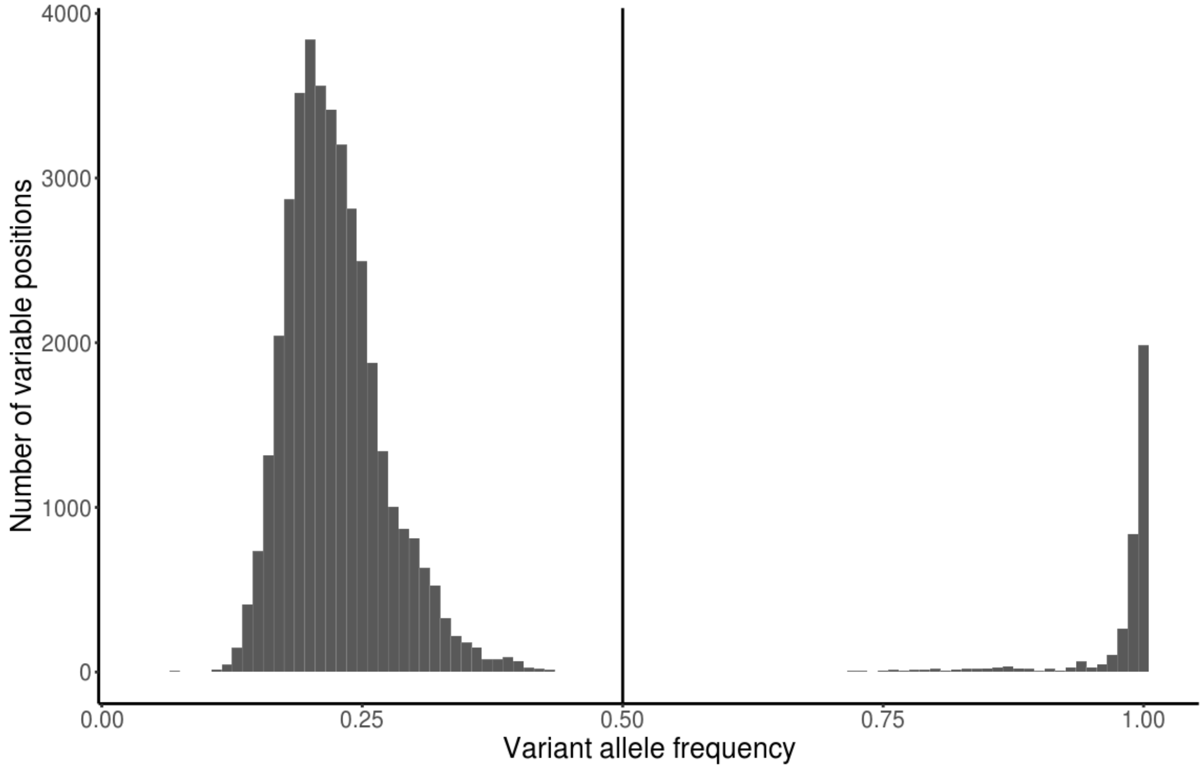

**Supplementary Fig. 8 | Distribution of variant allele frequencies based on 163 chloroplast genomes**

Variant allele frequency diagram indicating the frequency of reads supporting the variant allele aligned to a given locus. These reads might originate either from the chloroplast DNA or from mitochondrial/nuclear copies of specific chloroplast regions (mtptDNA/NUPTs). Therefore, genuine fixed mutations in chloroplasts can be represented by allele frequency lower than one. We observe two clearly separated peaks of allele frequency due to significant prevalence of chloroplast DNA. The left peak is likely to represent loci mutated in mitochondrial (or nuclear) copies of the specific chloroplast regions, but not on the chloroplasts themselves. The right peak thus is likely to represent genuine chloroplast fixed mutations. We only focused on the chloroplast fixed variant alleles by setting a threshold of >0.5.

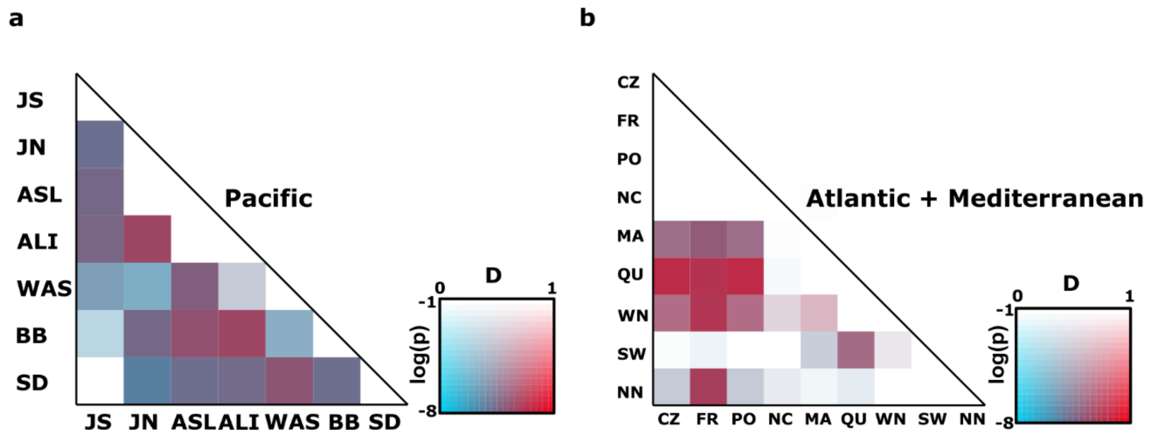

**Supplementary Fig. 9 | Matrix depicting Patterson's D-statistic (aka ABBA-BABA-statistics) for Pacific and Atlantic populations separately.**
Red colors indicate higher D-values, and more saturated colors towards the lower edge of the color legend indicate greater statistical significance (log(p)). Significant and high D-values indicate admixture between the two populations, but the direction of gene flow is not estimated. A signal of admixture can be caused by direct gene flow between the two populations or by genetic input from into both from the same "ghost" (=unsampled) population.

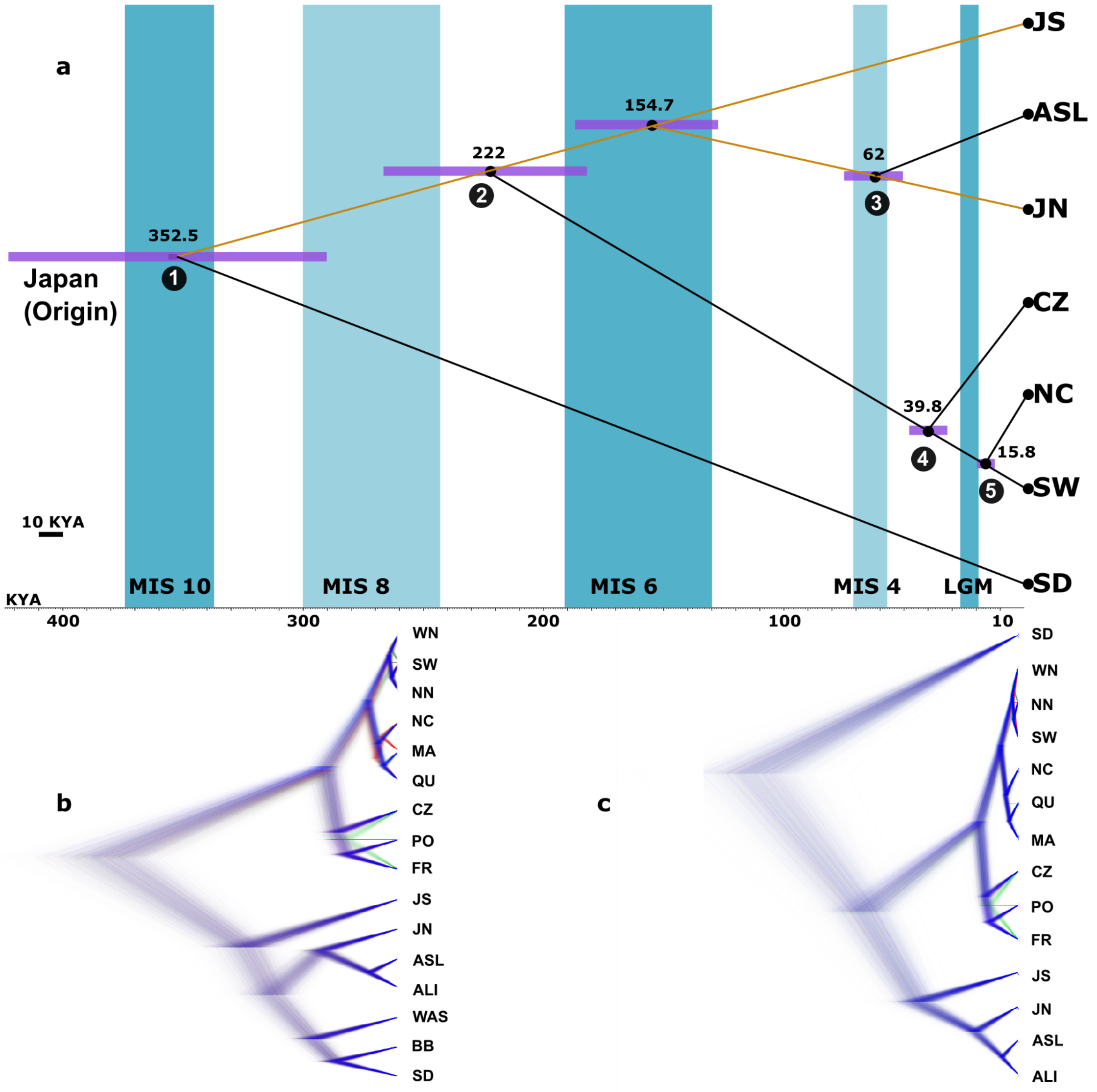

**Supplementary Fig. 10 | Coalescent phylogeny using the multi-species coalescent based on reduced data set of seven populations without any admixture**

**a**, Tree topology based on SNAPP analysis using a reduced data set comprising seven populations, excluding any populations involved in gene flow /admixture (D-statistics, Supplementary Fig. 9). Blue bars indicate glacial periods with Marine Isotope Stages (MIS) alternating with warm to cool interglacial periods with Last Glacial Maximum (LGM, 26.5–19 kya) at the right hand side. Purple bars across nodes indicate the 95% Highest Posterior Density (HPD) of the estimated divergence time. Key divergence events 1–5 are shown as in Fig. 4,6. Major splits 1-5 have largely similar age estimates compared to Fig. 4 main manuscript. **b**, DensiTree cloudogram with all 16 populations, thus not accounting for admixture and creating error in the placement of SD. **c**, Cloudogram with the most strongly admixed populations WAS and BB, removed.

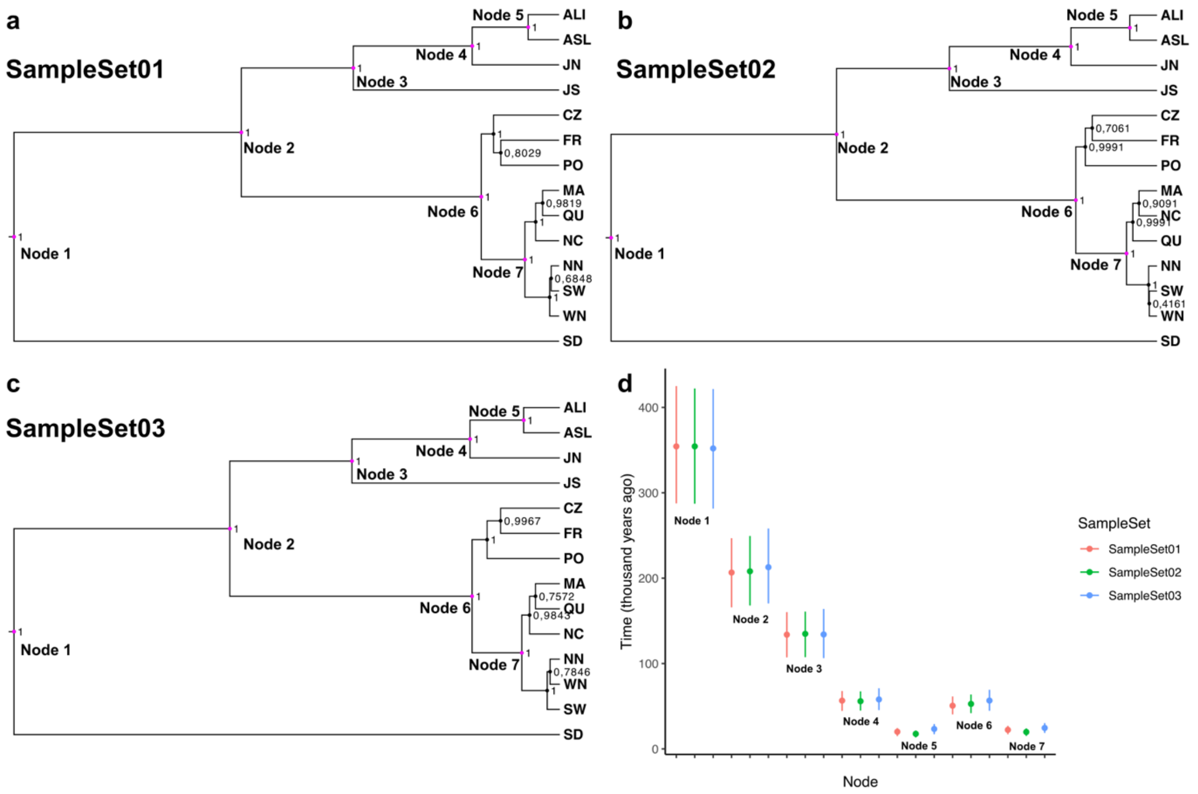

#### Supplementary Fig. 11 | Replicate coalescent phylogenies based on different replicate genotype sets per population.

**a-c,** Tree topologies for the three independent runs based on different sample sets using SNAPP are depicted. Major splits numbered 1–7 show a consistent topology across the three independent runs. Note the uncertainties regarding the topologies within the recent splits in the Northwest Atlantic, Northeast Atlantic, and Mediterranean Sea, respectively. **d**, Dating estimates and 95% highest posterior density (HPD) interval for the 7 consistent nodes for three different sample sets using SNAPP.

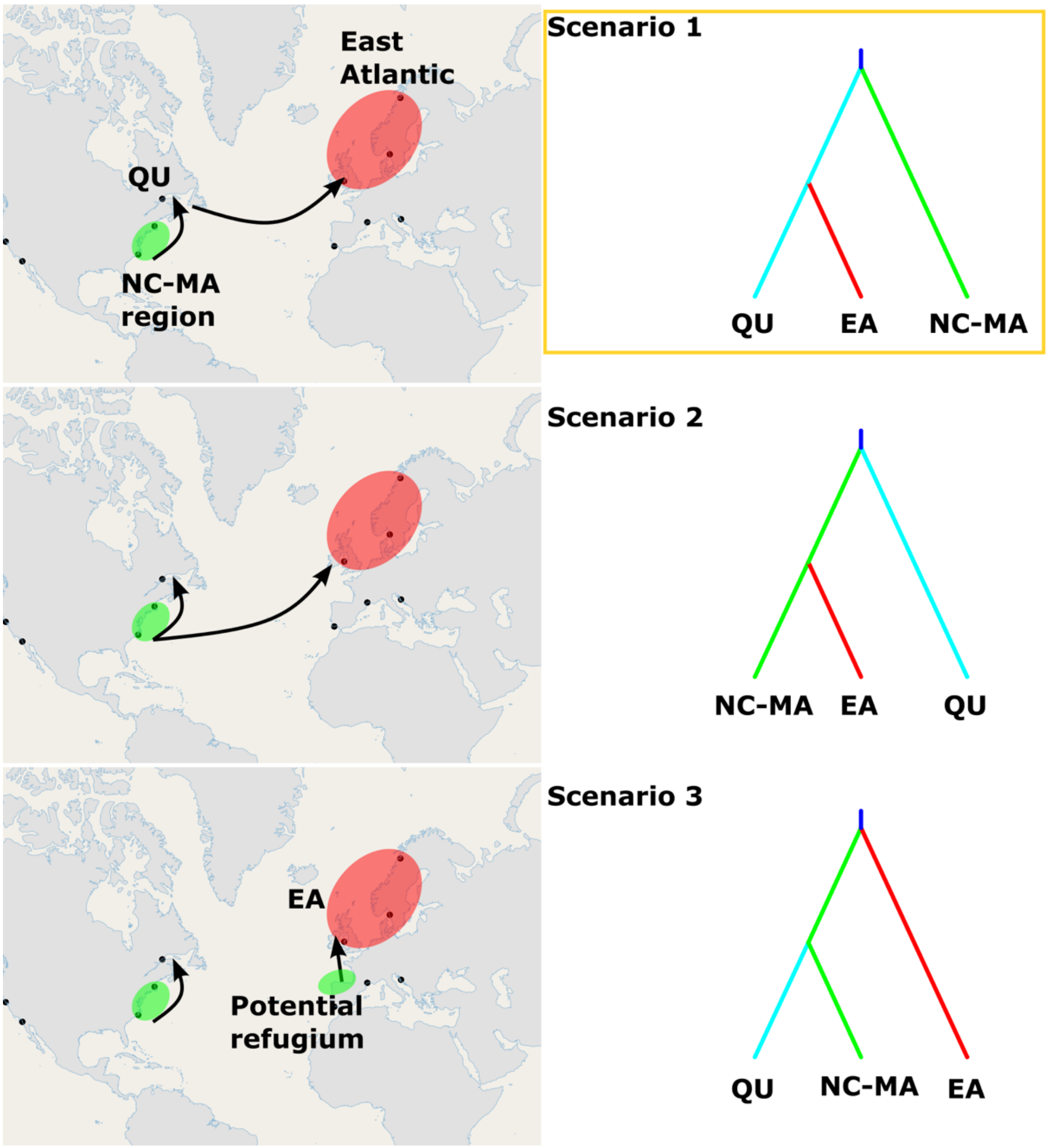

**Supplementary Fig. 12 | Most likely recolonization scenario of the Atlantic Ocean after the Last Glacial Maximum (LGM) using approximate Bayesian computation (ABC).**

Genetic diversity (Fig 1b,c), the coalescent tree (Fig. 4) and chloroplast haplotype network (Fig. 2g) indicate that the NC-MA region was one of the glacial refugia during the LGM, with a separate refugium in the Mediterranean. Southern Brittany, France has been proposed as a refuge for rocky intertidal/subtidal species (Jenkins et al., 2018, Maggs et al 2008), but a site in Brittany was not included in the present study. Approximate Bayesian computation implemented in the package DIYABC (Collin et a. 2021) was used to test the possible recolonization pathways in the Atlantic Ocean after the LGM. **Scenario 1**: the NC-MA region recolonized QU as stepping stone and subsequently the Northeast Atlantic. **Scenario 2**: NC-MA region recolonized QU and the Northeast Atlantic in parallel (no stepping stone). **Scenario 3**: NC-MA region only recolonized the northern NW Atlantic (QU), while a different early diverging and non-sampled refuge recolonized the Northwest Atlantic. **Scenario 1** was found to be the most likely scenario.

Collin F-D, Durif G, Raynal L, Lombaert E, Gautier M, Vitalis R, Marin J-M, Estoup A (2021) Extending approximate Bayesian computation with supervised machine learning to infer demographic history from genetic polymorphisms using DIYABC Random Forest. Molecular Ecology Resources 21: 2598-2613 doi https://doi.org/10.1111/1755-0998.13413
Jenkins T, Castilho R, Stevens J (2018) Meta-analysis of northeast Atlantic marine taxa shows contrasting phylogeographic patterns following post-LGM expansions. PeerJ 6: e5684
Maggs CA, Castilho R, Foltz D, Henzler C, Jolly MT, Kelly J, Olsen J, Perez KE, Stam W, Väinölä R, Viard F, Wares J (2008) Evaluating signals of glacial refugia for North Atlantic benthic taxa Ecology 89: S108-S122 doi https://doi.org/10.1890/08-0257.1

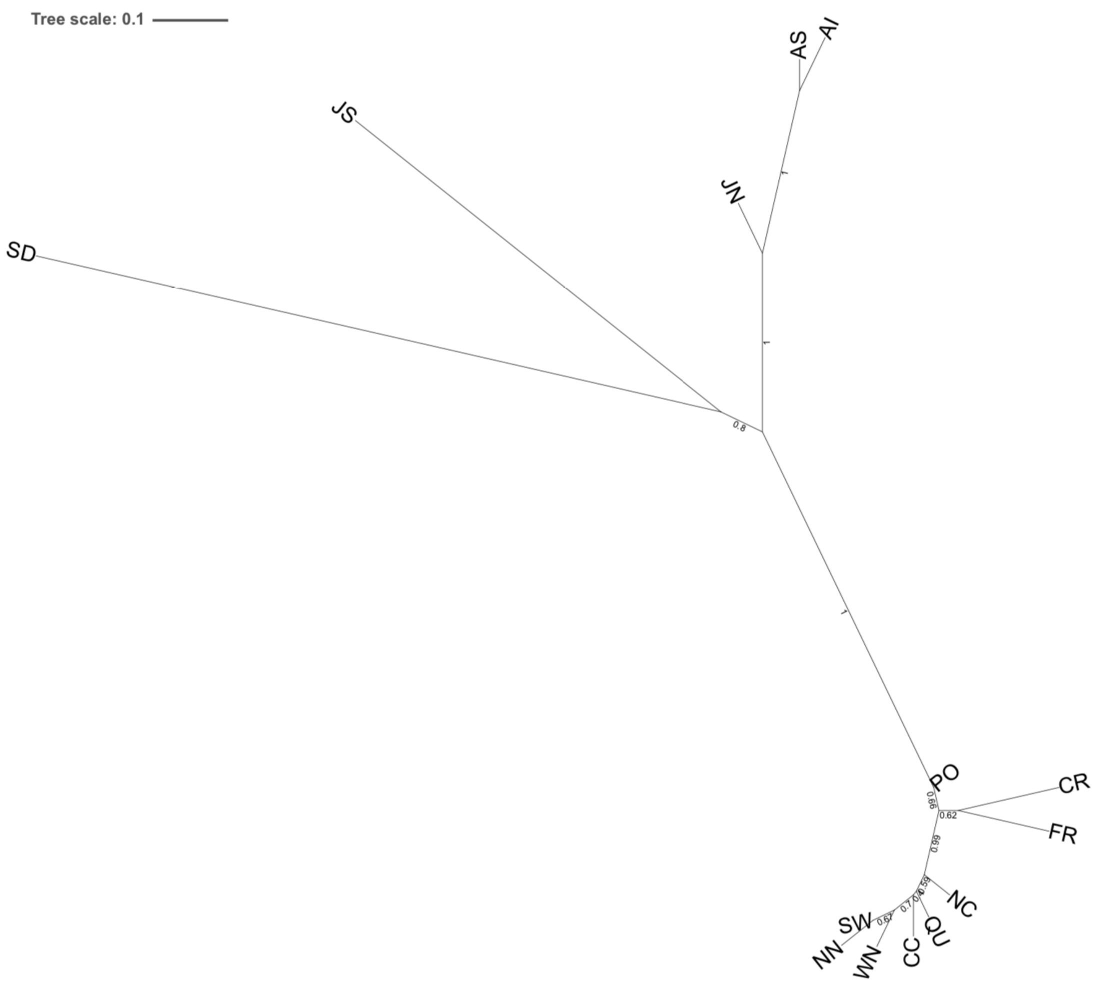

**Supplementary Fig. 13 | Unrooted phylogenetic tree using ASTRAL**
The data set includes the same 14 populations as in Fig. 4, and Supplementary Figures 10, 11. Heavily admixed populations (BB, WAS) were excluded. The final dataset for consideration included 129 samples and 20,100 aligned transcripts. Of the 18,311 core genes that were present in 97.9% of all samples, a random subset of 617 genes were selected. CDS and protein sequences from each transcript alignment were predicted using GeMoMa (v1.7). Gene trees were constructed by first aligning CDS sequences together using MAFFT (v7.475). (parameters: mafft --localpair --phylipout --maxiterate 1000), then generating individual gene trees with IQTREE (v2.1.2) (parameters -B 1000 -m K2P -T auto). With all estimated gene trees as input, the population tree was estimated with ASTRAL v5.7.3, using a map file to join multiple samples representing the same population. For further details and references see Supplementary Note 3.

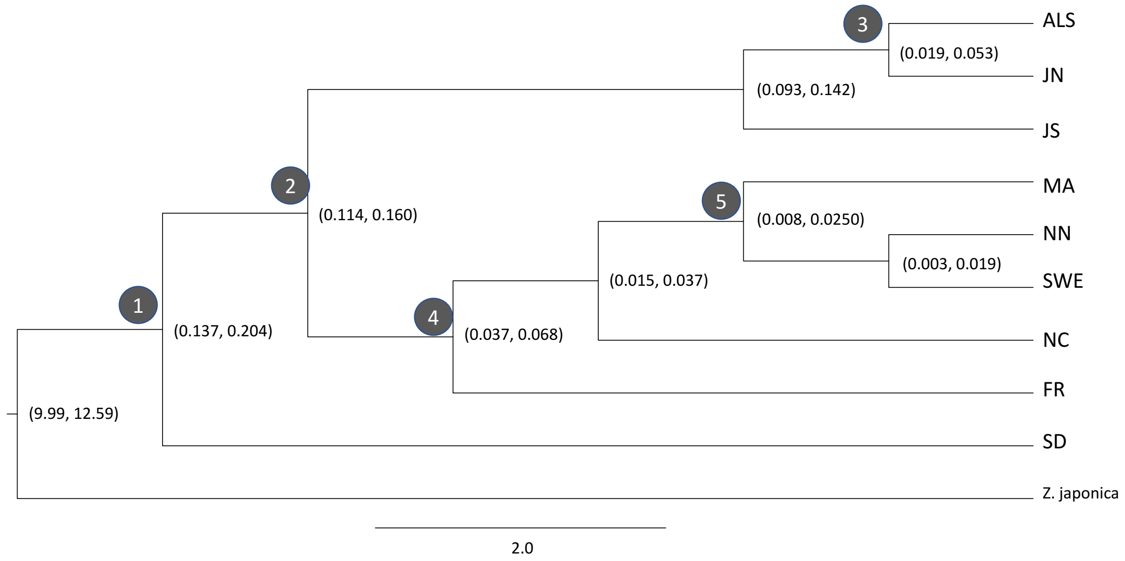

#### Supplementary Fig. 14 | starBEAST phylogenetic tree including divergence time estimation

Five major divergence /colonization events are numbered 1...5 as in Figs. 4, 6, Supplementary Figure 10. The XML file was generated using BEAUTi from 70 randomly selected alignments from a subset of 4 samples per population. Parameters used in the StarBEAST2 analysis were: gene ploidy = 2; constant population sizes; population size parameter = 0.03 (assuming $N_e$ = 10,000 and generation time 3 yrs); Gamma site model; estimated substitution rate; HKY substitution model; with estimated kappa parameter; empirical nucleotide frequencies; strict molecular clock; estimated clock rate; Yule model; Outgroup=*Z. japonica*; outgroup divergence time constrained with a lognormal prior [M=11.01; S=0.01; mean in real space, use originate]; MCMC chain length = 200,000,000; store every 200,000; pre-burnin = 0. StarBEAST2 was run in triplicate, inspecting each run with Tracer to check for model convergence (20% burn-in; Effective sample sizes [ESS]>300). TreeAnnotator was used to summarize each run (20% burn-in, median peak height, 0.5 posterior probability limit) which were then combined using LogCombiner. For further details and references see Supplementary Note 3.

### Supplementary Data

**Supplementary Data 1: Data coverage (.xls)**
**Supplementary Data 2: Mapping rate (.xls)**
**Supplementary Data 3: Genbank accession numbers (.xls)**
